# Reversing protonation of weakly basic drugs greatly enhances intracellular diffusion and decreases lysosomal sequestration

**DOI:** 10.1101/2023.04.19.537456

**Authors:** Debabrata Dey, Shir Marciano, Anna Poryvai, Ondřej Groborz, Lucie Wohlrábová, Tomás Slanina, Gideon Schreiber

**Affiliations:** Department of Biomolecular Sciences, Weizmann Institute of Science, Israel; Institute of Organic Chemistry and Biochemistry of the Czech Academy of Sciences, Flemingovo náměstí 542/2, Prague 6, 160 00 Czech Republic

**Keywords:** Small molecule drugs, diffusion, in-cell, lysosome

## Abstract

For drugs to be active they have to reach their targets. Within cells this requires crossing the cell membrane, and then free diffusion, distribution, and availability. Here, we explored the in-cell diffusion rates and distribution of a series of small molecular fluorescent drugs, in comparison to proteins, by microscopy and fluorescence recovery after photobleaching (FRAP). While all proteins diffused freely, we found a strong correlation between p*K*_a_ and the intracellular diffusion and distribution of small molecule drugs. Weakly basic, small-molecule drugs displayed lower fractional recovery after photobleaching and 10-to-20-fold slower diffusion rates in cells than in aqueous solutions. As, more than half of pharmaceutical drugs are weakly basic, they, are protonated in the cell cytoplasm. Protonation, facilitates the formation of membrane impermeable ionic form of the weak base small molecules. This results in ion trapping, further reducing diffusion rates of weakly basic small molecule drugs under macromolecular crowding conditions where other nonspecific interactions become more relevant and dominant. Our imaging studies showed that acidic organelles, particularly the lysosome, captured these molecules. Surprisingly, blocking lysosomal import only slightly increased diffusion rates and fractional recovery. Conversely, blocking protonation by *N-*acetylated analogues, greatly enhanced their diffusion and fractional recovery after FRAP. Based on these results, *N*-acetylation of small molecule drugs may improve the intracellular availability and distribution of weakly basic, small molecule drugs within cells.

## INTRODUCTION

Drug molecules alter many biochemical reaction pathways inside the cell by interacting with proteins, DNAs, RNAs, or others. By this, they influence one or more specific cellular activities (Ong et al., 2009; Schreiber, 2005; Stockwell, 2004). Due to their bioactivity, stability, rather simple chemical synthesis allowing for industrial bulk scale production and the easiness of administration, most drugs are small molecules (< 1 kDa) (Ngo and Garneau-Tsodikova, 2018; Veber et al., 2002). Over half of these drugs have a significant degree of hydrophobicity and are weakly basic at intracellular pH of 7.4 (Charifson and Walters, 2014; Lipinski et al., 1997). Hydrophobicity is required to cross the lipid bilayer of the cell membranes, and polar functional groups solubilize the drug in the cellular medium (Lipinski et al., 1997). The balance between p*K*_a_ and logP (partition coefficient of small molecules as measured from their partition between octanol/water) is critical for successful drug design (Manallack, 2007; Meanwell, 2011). Lipinski’s “Rules of 5” suggests that logP should be between 1 and 6 to be a candidate for successful oral administration. Small molecule drugs can be acidic or basic depending on their p*K*_a_ values (Lipinski et al., 1997). Drugs with p*K*_a_ ∼ 0-7 are considered acidic, and p*K*_a_ ∼ 8-14 are regarded as basic or alkaline. Basic drugs with p*K*a ˃8 are more common than their acidic counterparts (p*K*a ˂ 7, ∼60:40 ratios). In some cases, lowering their p*K*_a_ to the range between 6 and 7, has proven beneficial (Charifson and Walters, 2014). For a long time, mechanisms such as passive diffusion (Di et al., 2012) and carrier-mediated drug delivery (Dobson and Kell, 2008) or a combination of both (Sugano et al., 2010) have been used to explain intracellular drug delivery. However, transport of a drug across the cellular membrane is only one part of a successful delivery. Intracellular diffusion and bio-distribution needs to be taken in account when assessing the drug mode of action.

In previous work, we studied the diffusion of fluorescent drug molecules in crowded environments *in vitro* (Dey et al., 2022). However, the cellular cytoplasm of eukaryotic cells is by far more complex (Model et al., 2021). Intracellular membranes, multiple organelles (Ellis, 2001; Fulton, 1982) and a large diversity of macromolecules, in terms of size, charge and hydrophobicity, render the cellular interior extremely complex and heterogeneous (Milo and Phillips, 2015; Neurohr and Amon, 2020). Moreover, the pH inside cellular organelles can vastly differ from the cytosolic pH (Madshus, 1988). Organelles like lysosomes, late endosomes, Golgi, and secretory vesicles are acidic and have been known for sequestering weakly basic drugs (Asokan and Cho, 2002; Proksch, 2018). Therefore, predicting diffusion of a particular drug molecule in the dense cytoplasm is a complex task and requires sub-organelle resolution. For example, positively charged (basic) drugs designed to target nuclear DNA may bind cytoplasmic matter through nonspecific interactions (Long et al., 2022; Tiwari et al., 2018), they could be sequestrated within the lysosome, or diffuse slower than expected due to crowding and nonspecific interactions (Gotink et al., 2011; Jansen et al., 1999; Zhitomirsky and Assaraf, 2015).

The mobility of a drug in the cellular interior has been shown to be a good proxy for intracellular drug delivery and other processes, such as nonspecific binding or aggregation (Dey et al., 2022). The primary tool for measuring small molecule diffusion inside the cell is fluorescence recovery after photobleaching (FRAP), which inherently requires fluorescent drug molecules. FRAP with a confocal microscope is a sufficiently sensitive, versatile, and easy-to-use method suitable for measuring the diffusion of weakly fluorescent molecules. Unfortunately, single-molecule fluorescence correlation spectroscopy (FCS), a gold-standard method for measuring diffusion of fluorescent particles, cannot be applied for most small molecule drugs due to their weak fluorescence (Dey et al., 2022, 2021; Zotter et al., 2017).

In this work, we determined the intracellular diffusion coefficients of a series of fluorescent dyes and drugs compared to protein diffusion by FRAP (Figure 1 and Table 1). Specifically, we used Line-FRAP, which has a much higher time resolution and therefore can measure faster diffusion rates in comparison to standard FRAP (800 Hz versus 50 Hz, for detailed explanations, see (Dey et al., 2022) (Dey et al., 2021)). A pictorial representation of Line FRAP performed in a live HeLa cell treated with a fluorescent drug is shown in Figure 2A. The small red circle denotes the bleached area which is done by using a Clip tornado (of 2x2 pixel size). The white line indicates the scanning region which was performed along at a rate of 800 Hz in order to record the FRAP as a function of time. The Line-FRAP cross section profile as a function of time is shown in Figure 2B. The vertical direction of the line denotes the time progress. The vertically marked white rectangular Region of Interest (ROI), in the left panel (Figure 2B) shows the fluorescence intensity recovery over time, which is plotted in Figure 2C. The horizontal rectangular region of interest (ROI) outlined in white (in the right panel of Figure 2B) denotes the time frame immediately after the photo-bleach pulse (the highly intense green is the bleach pulse). The time interval between each line scanning measurement is ∼1.25 milliseconds. This horizontal white ROI generates the fluorescence intensity over the scanning line distance which is plotted in Figure 2D. This profile gives the effective bleach radius when fitted with a gaussian function (Dey et al., 2022, 2021). The apparent diffusion coefficients derived from FRAP measurements are termed D_confocal_ are calculated using equation 1:

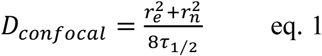

**Figure 1.**
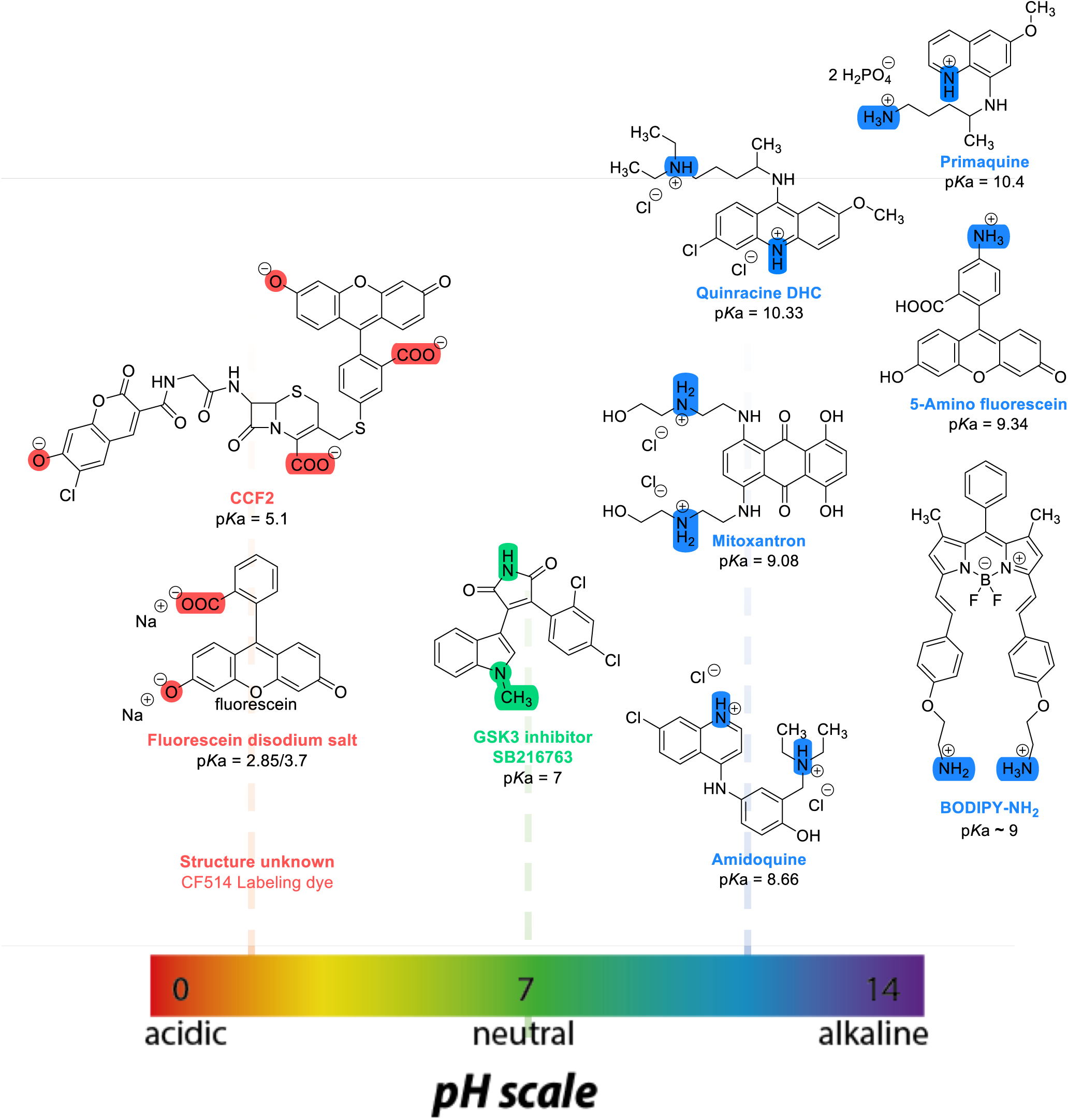
Chemical structures and p*K*a values (prediction from ChemAxon software) of the small molecules and drugs used in this study.

**Figure 2.**
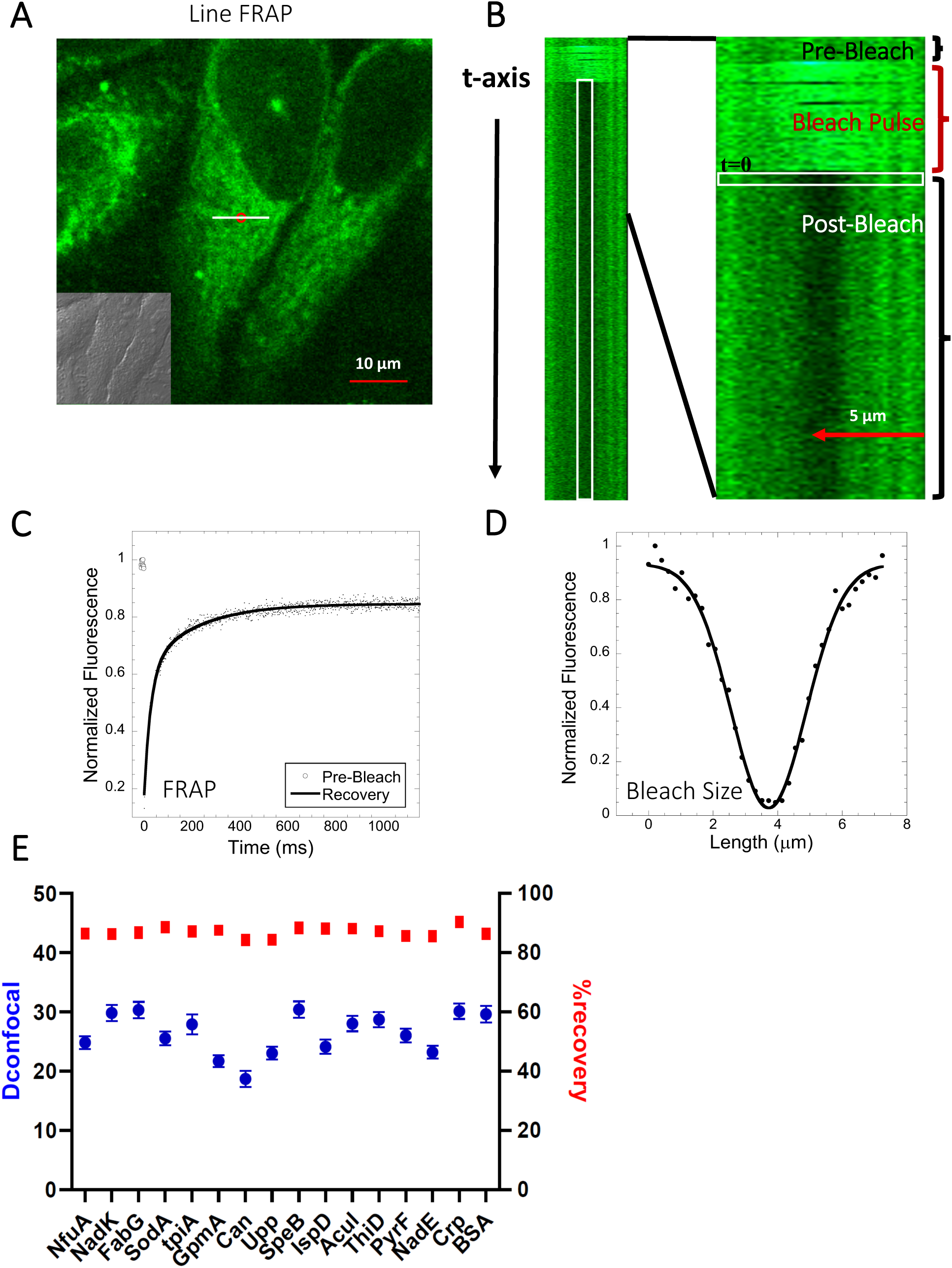
Schematic representation of Line-FRAP & Diffusion coefficients of proteins in the HeLa cells. (A) Line-FRAP performed in a fluorescent drug labeled HeLa cell. The white line represents the scanning line; The red circular area denotes the bleaching area. (B): A single scanning line as a function of time (in the vertical direction) including the bleach is shown. The white vertical rectangular region of interest (ROI), which was used for analysis. A close-up view of the Line-FRAP profile, is also shown where the horizontal rectangular area (white colored marks ROI) denotes the fluorescence intensity as a function of length across the scanning line. (C) Average of 30 FRAP recovery curves as a function of time (N=30; Correlation coefficient of exponential fit: R=0.98) and (D) Averaged bleach size profile with gaussian fits (N=30; R=0.99) are shown. (E) Comparison of diffusion coefficients (D_confocal_ in blue circles) and percentage of recoveries (in red squares) of bacterial proteins and BSA as measured in HeLa cell cytoplasm are shown. Error bars represent SE calculated from fitting the FRAP progression curves, which are averaged over at least 30 independent measurements.

**Table 1:**
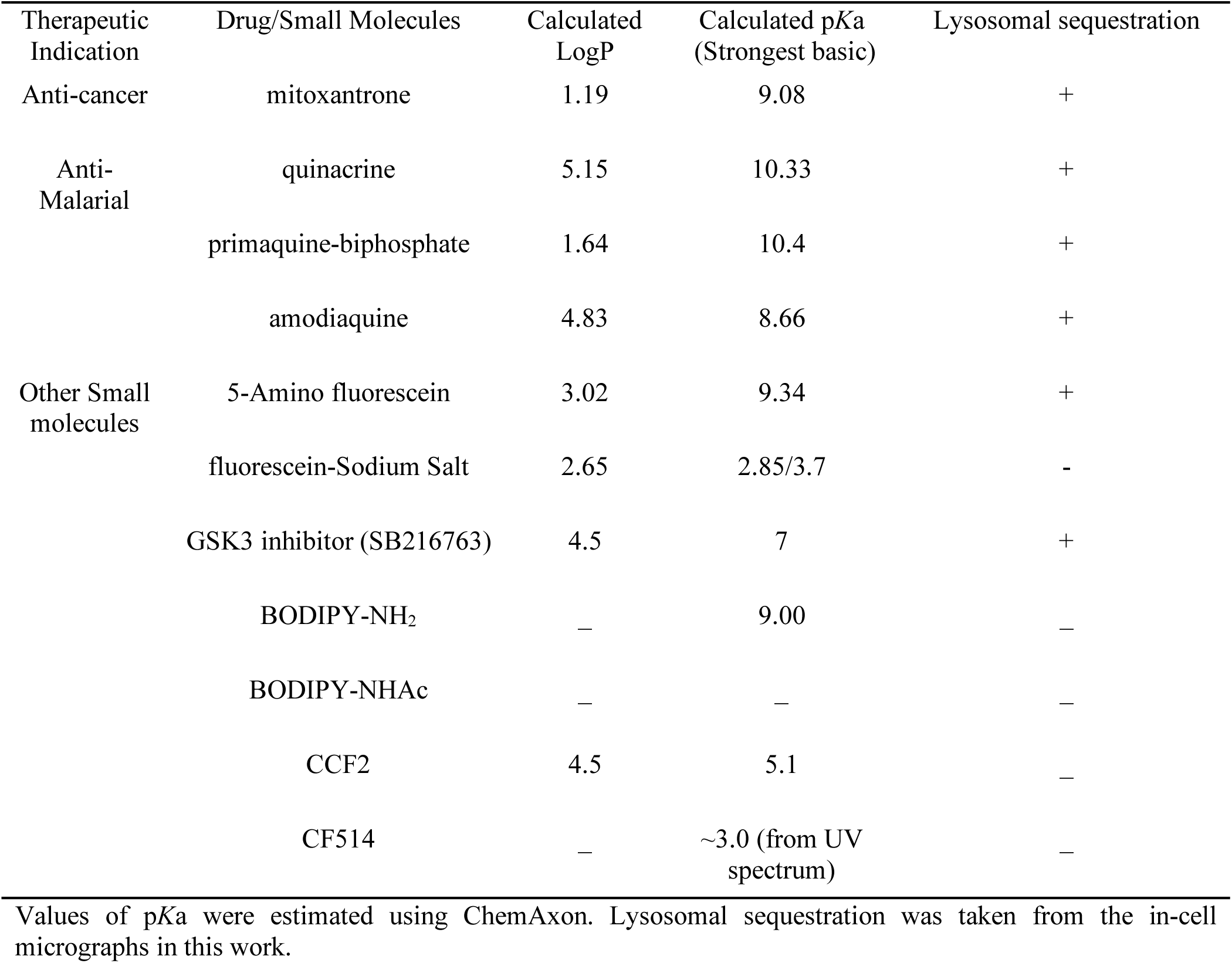
LogP, p*K*a and lysosomal sequestration of drugs used in this study (Predicted by ChemAxon software, and obtained from drug bank data base (Zhitomirsky and Assaraf, 2015))

where t1/2 is the half-time of recovery and re and rn are the effective and nominal bleach radii values (for details see (Dey et al., 2021)). D_confocal_ values for proteins determined by Line-FRAP agreed with the diffusion rates measured by FCS (Dey et al., 2021). For small molecules, Line-FRAP only provides relative 3D diffusion coefficients (Dey et al., 2022). We studied the following fluorescent drugs: the antimalarial drugs quinacrine dihydrochloride (Ehsanian et al., 2011), primaquine phosphate (Camarda et al., 2019) and amidoquine (Olliaro and Mussano, 2009). The anti-cancer drug mitoxantrone (Shenkenberg and Von Hoff, 1986), and the glycogen synthase kinase GSK3 inhibitor SB216763 (Wagman et al., 2005). In addition, we monitored the diffusion of the diagnostic staining agent fluorescein disodium salt (Jampol and Cunha-Vaz, 1984), the fluorogenic substrate CCF2 (Zuverink and Barbieri, 2015) and boron dipyrromethene (BODIPY) analogues, which are used as fluorescent markers for lipids, membranes, and other lipophilic compounds (Poryvai et al., 2022).

The diffusion coefficients and their cellular distributions showed very slow diffusion of the basic compounds, i.e., low FRAP recovery, combined with an accumulation of the molecules in lysosomes. Surprisingly, while inhibiting the V-ATPase H^+^ pump using Bafilomycin A1 (Spugnini et al., 2010; Wang et al., 2021) or sodium azide (Hiruma et al., 2007) inhibited accumulation in lysosomes, cellular diffusion and fractional recovery increased only slightly. Contrary, blocking protonatable amino groups by acetylation greatly enhanced diffusion and FRAP recovery of studied small molecules. As over half the small molecule drugs are basic, our findings should be considered in drug assessment and development, as slow diffusion and low FRAP recovery is directly related to reduced cellular activity. Contrary to small molecules, we show here that diffusion rates within the cytoplasm of HeLa cells of 16 different *E.coli* proteins with p*K*a values of 4.5-8, are within the expected range, suggesting that hindered diffusion is mainly a problem of small molecules.

## RESULTS

### Acidic Proteins diffuse freely in the HeLa cytoplasm

We have previously shown that diffusion in the cellular cytoplasm of E-fts and baeR, two bacterial globular proteins, aligns with the expected values considering the ∼2-3-fold higher macro-viscosity (Dey et al., 2021). Here, we extended this study to include 16 additional proteins (mostly from *E. coli*) whose oligomeric states in solution were previously characterized (Table S1 and (Marciano et al., 2022)). The aim was to determine how different heterologous proteins (*E.coli* proteins in HeLa cells) diffuse. The proteins showed D_confocal_ values of 18-30 µm^2^s^-1^ in HeLa cell cytoplasm, which is typical for a cytoplasmic protein diffusion coefficient (Figure 2E). Moreover, full FRAP recovery was observed, indicating free diffusion. Table S1 provides the predicted isoelectric points and net charges at pH 7.4 of the purified proteins as calculated by Prot Pi, showing p*K*a values of 4.5-8, which are representative of the majority of *E.coli* cytosolic proteins. The results show that acidic and neutral proteins, even if not in their native environment, do not stick to the HeLa cell cytoplasm or membranes.

### Negatively charged small molecules rapidly diffuse in live cells

Fluorescein is a negatively charged small organic molecule (p*K*_a_’s = 2.85, 3.7, M.W. = 332 Da, Figure 1 and Table 1). We showed in previous work that fluorescein diffusion is slower in the presence of crowder proteins like BSA, lysozyme and, myoglobin, even at relatively low protein concentrations (Dey et al., 2022). Here, the Fluorescein dye was micro-injected in live HeLa cells, followed by FRAP measurements (Figure 3 A-D). We found that the diffusion coefficient of fluorescein in PBS (D_confocal_ of 56.5±2.4 µm^2^s^-1^) was reduced to D_confocal_ of 38.5±2.0 µm^2^s^-1^ in HeLa cytoplasm (Figure 3C). In the cell nucleus, a value of D_confocal_ of 43.2±2.2 µm^2^s^-1^ was determined, while in concentrated HeLa cell extract (equivalent protein concentration of ∼100 mg/mL), D_confocal_ of 49.6±1.9 µm^2^s^-1^ was measured. The faster D_confocal_ in cell extract compared to live-cell cytoplasm may result from the higher crowding density of the latter. The lowering of diffusion coefficients of negatively charged fluorescein correlates with the increasing crowding density of the medium moving from simple buffer to cells. For comparison, we also incubated HeLa cells for 24 hrs with fluorescein, which is sufficient time for it to accumulate in the cells by diffusion (Figure S1 A-B). While micrographs did not show dye aggregation inside the cell cytoplasm (Figure S1A-B) and diffusion rates are similar to those determined by microinjection (Figure S1C and S1D), the fractional recovery dropped from 0.95 to 0.65. This suggests that a fraction of fluorescein is immobile in the cell after 24 h incubation, and is not diffusing.

**Figure 3.**
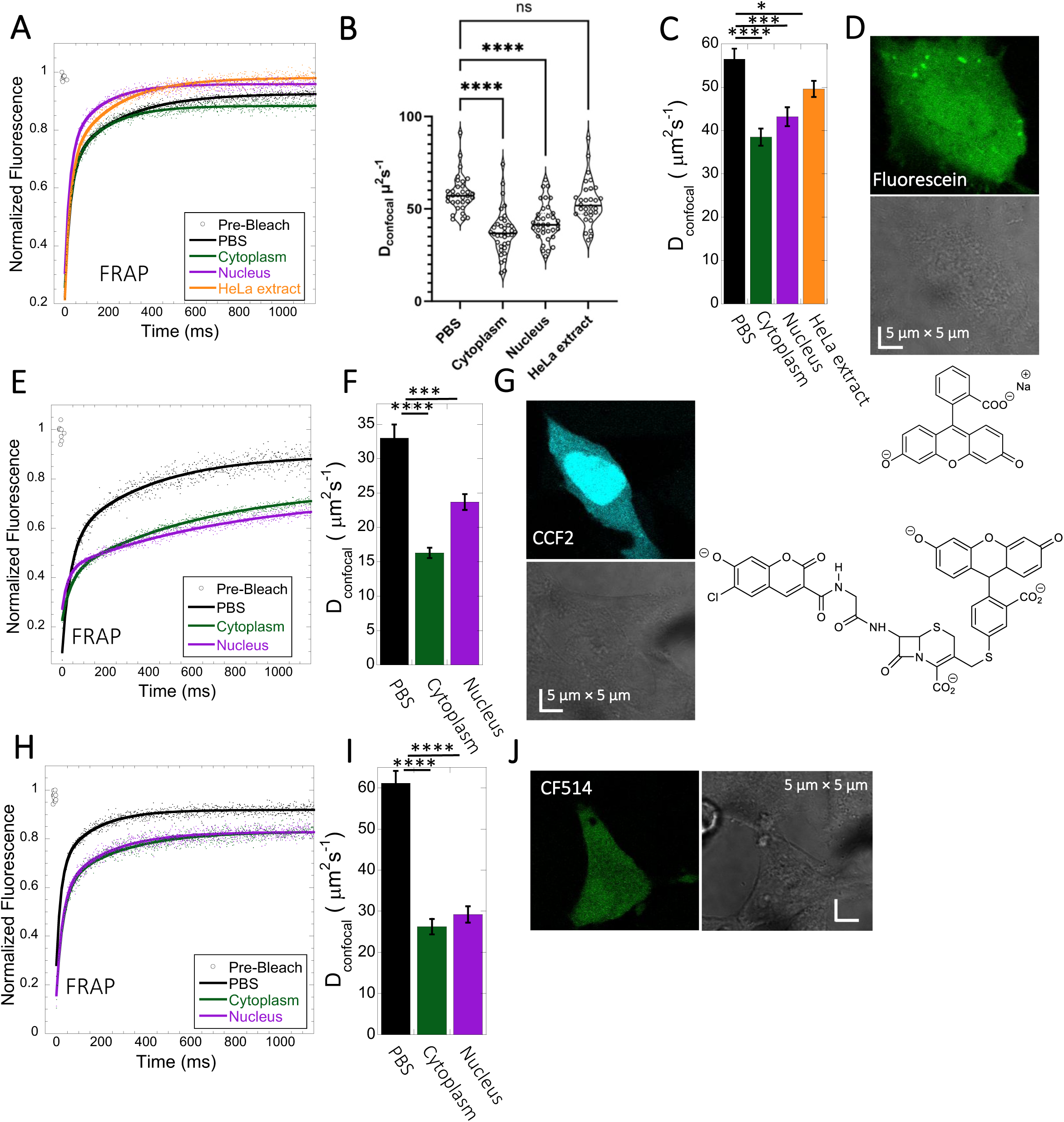
Fluorescein, CCF2 and CF514 diffusion in PBS and inside HeLa cells. Comparative merged FRAP profiles (N= 30; R= 0.99 for each of the fits) in PBS and inside HeLa cells with exponential fits are shown for (A) Fluorescein, (E) CCF2 and (H) CF514. Comparative D_confocal_ values for (C) Fluorescein (F) CCF2 and (I) CF514 are also shown. (B) D_confocal_ values as calculated from individual FRAP curves for Fluorescein (see also Fig. S2). D_confocal_ values as calculated from the merged FRAP curves shown in panel A. (D) HeLa cells after Micro-injection of Fluorescein; (G) CCF2; and (J) for CF514. Error bars represent SE calculated from fitting the FRAP progression curves of 30 merged independent measurements. Statistical significance calculations are detailed in the materials and methods section.

For calculating the D_confocal_ values shown in Figure 3C, 30 individual Line-FRAP curves were merged for signal to noise reduction (Figure 3A) (Dey et al., 2022, 2021). For comparison, we calculated D_confocal_ values from individual Line FRAP progression curves in HeLa cytoplasm (Figure S2A and S2B), obtaining the same mean value as for the merged curve (Figure S2B and S2D), however, with much larger variation over the mean. To estimate whether the large variation from the mean of the individual curves is due to different fluorescein concentrations in the individual measurements, we plotted the fluorescence signal (which is directly related to the concentration) versus the FRAP recovery rate (Figure S3A) and versus the percent recovery (Figure S3E). The lack of correlation suggests that the variation in FRAP rates of the individual measurements is due to the low signal to noise of the individual curves versus the average. Next, we calculated D_confocal_ also for fluorescein in PBS, nucleus and HeLa cell extract (Figure 3B). The mean values from individual curves are very similar to those calculated from the merged curves (Figure 3C), with a normal distribution of the individual calculated D_confocal_ values around the mean. Moreover, also for the values measured in the nucleus, no relation between fluorescein concentration and FRAP rates or percent recovery was observed (Figure S3B and F). Overall, this suggest that merged curves can be used to calculate D_confocal_ values, as they give much better signal to noise, with apparent minimal loss of information.

CCF2 is a fluorogenic substrate used for monitoring β-lactamase activity, with a p*K*a of ∼5.1 (Zlokarnik et al., 1998). It is composed of cephalosporin core linking B7-hydroxycoumarin to fluorescein (Figure 1). Its calculated Dconfocal values in PBS and inside cells are about half as fast as those measured for fluorescein (Figure 3E-G), and its fractional recovery is ∼0.7. No sign of aggregation is seen in micrographs inside the cell cytoplasm or nucleus (Figure 3G). The substantial slower rate of recovery, and lower fractional recovery of CCF2 relative to Fluorescein suggests its diffusion to be obstructed within the cell. This is in line with our prediction, that the observed substrate limited enzymatic degradation of CCF2 in HeLa cells is a consequence of its occlusion within the cellular cytoplasm (Zotter et al., 2017).

A third molecule for which we determined the diffusion coefficient is the labelling dye CF514. It is used to label proteins and has high quantum yield. As its structure has not been published, we estimated the p*K*_a_ value from the pH dependent UV-spectrophotometry measurements. The sharp changes of the UV-spectrum at pH∼3 (Figure S3I-J), as well as the redshift confirms a p*K*_a_ value of ∼ 3. The FRAP curves look similar to fluorescein (Figure 3H-J), with ∼90% recovery after FRAP. D_confocal_ in PBS was ∼ 61.1±3.0 µm^2^s^-1^, and 26.3±1.9 and 29.2±2.0 µm^2^s^-1^ in the cell cytoplasm and nucleus of HeLa cells, respectively (Figure 3I). The drop of D_confocal_ is attributed to cellular crowding. The results so far show that acidic small molecules diffuse well inside cells. Next, we focus on basic small molecules with higher p*K*a values.

### High FRAP recovery but slow diffusion measured for the glycogen synthase kinase (GSK3) inhibitor SB216763

The GSK3 inhibitor SB216763 (p*K*a=7.0, M.W.=371 Da, Figure 1) aggregates when dissolved in PBS buffer solution. However, in our previous study we have seen that aggregation of this particular small molecule drug can be reversed by adding BSA and the extent of de-aggregation depends on the amount of BSA added (Dey et al., 2022). Here, our aim is to compare diffusion behavior of the GSK3 inhibitor in solution phase to that found in live cell cytoplasm. Since the molecule has some inherent solubility issues in PBS buffer, we compared its diffusion also in BSA protein solution and in cell culture media DMEM (Dulbecco’s modified eagle medium). Comparative FRAP profiles and D_confocal_ rates of SB216763 in different mediums are shown in Figure 4A-C. The FRAP profiles and relative D_confocal_ values show very slow diffusion of SB216763 in HeLa cell cytoplasm of ∼0.61±0.03 µm^2^s^-1^ with a fraction recovery of 0.8 after 10s (Figure 4A-C). This is a 40-fold reduction in D_confocal_ compared to measurements in 100 mg/mL BSA protein solution, where SB216763 is not aggregating (D_confocal_∼24.3±0.5 µm^2^s^-1^ and fractional recovery of ∼0.9 - Figure 4A-C). In cell culture DMEM media (containing 5% serum protein) fractional recovery after FRAP is very low (Figure 4A), indicating that the drug is still aggregating (Dey et al., 2022). Interestingly, the D_confocal_ value in HeLa cell extract was higher ∼ 11.3±0.7 µm^2^s^-1^ as is its fractional recovery (0.9), showing that HeLa cell extract, like BSA is disaggregating SB216763. Co-localization with Lysotracker-633 (Figure 4D-E) shows that SB216763 colocalizes and is sequestered in acidic chambers of the cell lysosomes. Moreover, SB216763 accumulation into the lysosome increased over time (Figure 4D and E, 45-minute and 2-hour treatment). Varying drug dosages resulted in D_confocal_ values of 0.42-0.61 µm^2^s^-1^, suggesting that the slow diffusion is not related to SB216763 concentration (Figure S4A-B). To validate D_confocal_ values determined from merged progression curves, we calculated them also using individual Line FRAP profiles (Figure 4C vs 4B), with both methods showing the same mean values. Individual FRAP progression curves are shown in Figure S2E, which have a much lower signal to noise ratio in HeLa cell cytoplasm, but result in a similar mean D_confocal_ value as determined from the merged curves (Figure 4A-C and S2E-H). There may be a slight decrease in FRAP rates and percent recovery at higher SB216763 concentrations as determined from the individual profiles (Figure S3C and G).

**Figure 4.**
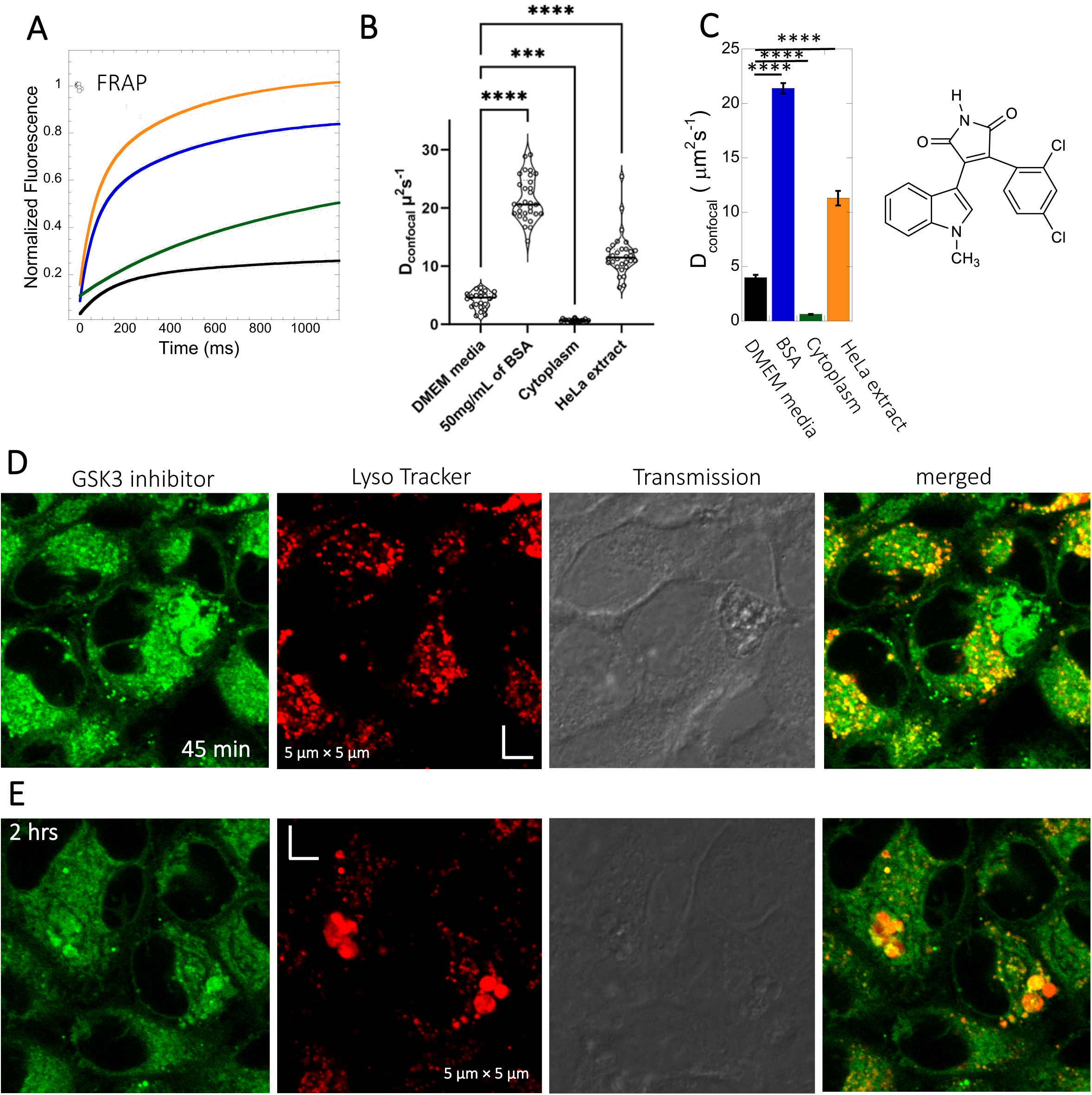
Diffusion of GSK3 inhibitor in PBS and inside HeLa cells. Comparative (A) averaged FRAP recovery profiles with exponential fits for GSK3 inhibitor (SB216763) (N= 30; R= 0.99 for each of the fits). Dots show prebleach fluorescence, Black line for DMEM, blue for PBS buffer + 50 mg/mL of BSA, green for HeLa cells and orange for HeLa extract. (B) D_confocal_ values calculated from individual FRAP curves. (C) D_confocal_ values calculated from merger of 30 independent FRAP curves. (D-E) Colocalization of GSK3 inhibitor in lysosomal compartments. HeLa cells were treated with Lyso Tracker 633 for 30 mins. GSK3 inhibitor treatment for (D) 45 mins and (E) 2 hours respectively at 10 µM concentrations are also shown. Error bars represent SE calculated from fitting the FRAP progression curves, which are averaged over at least 30 independent measurements. Statistical significance calculations are detailed in the materials and methods section.

### Despite its high solubility in aqueous solution, Quinacrine diffusion in HeLa cells is slow

Quinacrine DHC (M.W.= 472.9 Da, Figure 1) is a compound highly soluble in PBS (Dey et al., 2022) with a fast D_confocal_ rate of 41.9±3.0 µm^2^s^-1^ (Figure 5A-C). This rate dropped to D_confocal_∼2.5±0.1 µm^2^s^-1^ in HeLa cell cytoplasm, with fraction recovery of ∼0.8, in comparison to one in buffer. In HeLa cell extract a D_confocal_ rate of 15.1±0.6 µm^2^s^-1^ was determined, which lays between PBS and HeLa cytoplasm. D_confocal_ estimated from individual Line FRAP profiles and the merged Line FRAP profile (Figure 5B vs 5C) are also shown. Figure S2I-L shows that the merged FRAP profile yields less noisy data and gives a better fit, yet the mean D_confocal_ values are the same. As with SB216763, lower fraction recovery is observed at high quinacrine concentrations, but the FRAP rates seem to be concentration independent (Figure S3D and H). Micrographs of live HeLa cells show that the quinacrine DHC molecules aggregate in the cytoplasm. Co-localization with Lysotracker-633 (Figure 5D-E) show that quinacrine colocalizes and is sequestered in acidic chambers of the cell lysosomes. Moreover, quinacrine aggregation into the lysosome increased over time (Figure 5D and E, 45-minute and 2 hours treatment). Therefore, we measured diffusion instantaneously following micro-injection, and compared it to 2 and 24 hours of incubation (Figure S5A-B). Quinacrine accumulation in the lysosome was observed also immediately after micro-injection, with aggregation increasing over time. D_confocal_ of 4.2±0.2 µm^2^s^-1^ was calculated from line-FRAP immediately after micro-injection, slowing to 2.2±0.1 µm^2^s^-1^ following 2 hours incubations, with fractional recoveries of 0.63 and 0.57 respectively. After 24 hours of incubation the fractional recovery is less than 0.25, and thus the diffusion value is meaningless (Figure S5C and D). Next, we compared FRAP of the cells treated with 2 and 6 µM quinacrine solutions after 2- and 24-hours incubations (Figure S6A-F). The concentration of quinacrine had only a marginal effect on the results. Again, following 24 hours of incubation, most of the quinacrine is in the lysosomes.

**Figure 5.**
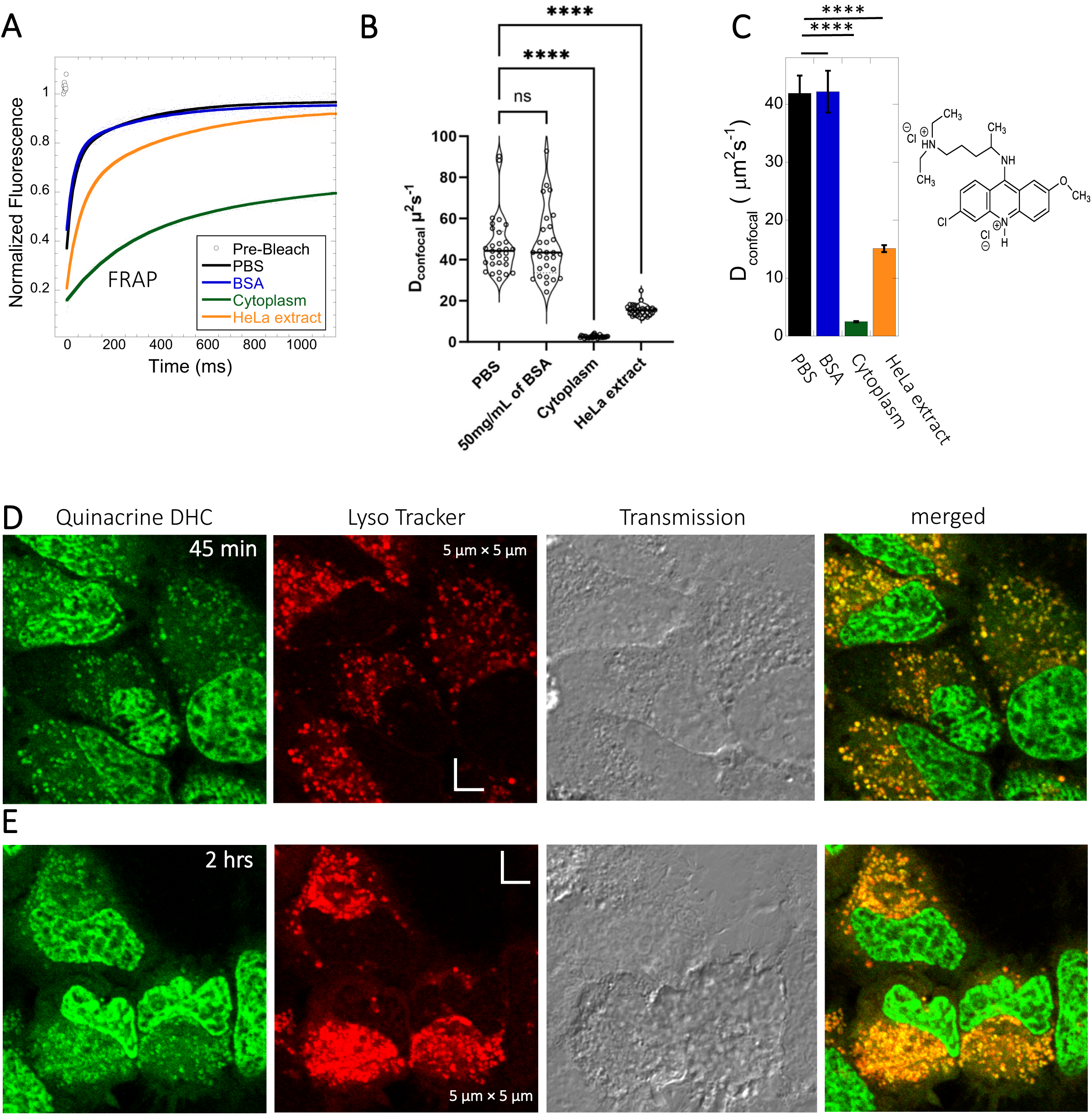
Diffusion of Quinacrine in PBS and in HeLa cells. Comparative (A) averaged FRAP recovery profiles with exponential fits for Quinacrine dihydrochloride (N= 30; R= 0.99 for each of the fits), (B) D_confocal_ values calculated from individual FRAP curves. (C) D_confocal_ values calculated from merger of 30 independent FRAP curves measured in PBS buffer, 50mg/mL of BSA, HeLa cell extract and in live HeLa cells. (D-E) Colocalization of Quinacrine DHC in lysosomal compartments are shown. HeLa cells were treated with Lyso Tracker 633 for 30 mins. Quinacrine DHC treatment was done for (D) 45 mins and (E) 2 hours respectively at 10 µM concentration. Error bars represent SE calculated from fitting the FRAP progression curves, which are averaged over at least 30 independent measurements. Statistical significance calculations are detailed in the materials and methods section.

### Mitoxantrone, primaquine and amidoquine all aggregate within cells

Mitoxantrone (anti-cancer) and primaquine phosphate (antimalarial) fluorescence with low quantum yields and weak photo-stability. However, due to their slow diffusion, the temporal resolution of standard XY-FRAP was sufficient for FRAP measurements (Figure 6A-H). Figures 6A and 6E show very low fraction recovery for these two drugs, suggesting that these weakly basic drugs are also sequestered in the cellular cytoplasm, as indeed shown in Figures 6B and 6F. Colocalization with LysoTracker red-633 show that Primaquine (S7A-B) and Amidoquine (S7C-D) are sequestered in the lysosomes. For amidoquine unfortunately, we could not collect any FRAP profiles as the dye did not survive the photobleached pulse. While we selected bleach spots to be small and located outside of lysosomes, this does not assure that some of the bleached area does not include smaller lysosomes. Therefore we investigated whether inhibiting lysosomal trapping will eliminate slow diffusion of cationic drugs.

**Figure 6.**
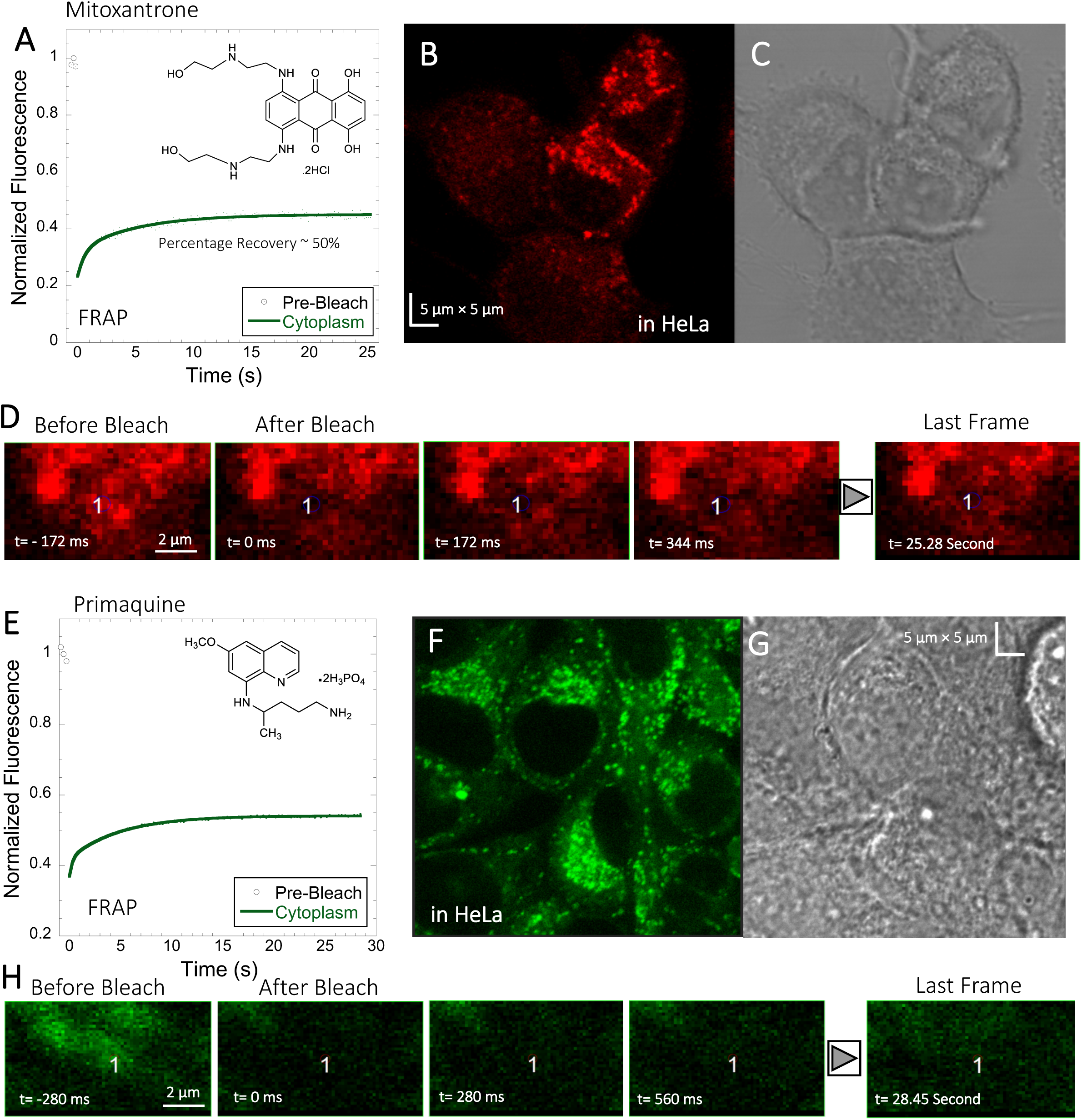
Mitoxantrone and Primaquine diffusion inside HeLa cells. Averaged XY-FRAP recovery profiles with exponential fits (N= 20; R= 0.98 for each of the fits) for (A) Mitoxantrone and (E) Primaquine in HeLa cells. Fluorescent and Transmission channel images of HeLa cells after 45 mins incubations with (B-C) Mitoxantrone and (F-G) Primaquine are shown. Time lapse micrographs of a portion of HeLa cell going through a classical rectangular XY-FRAP protocol for (D) Mitoxatrone and, (H) Primaquine are also shown.

### Inhibition of lysosomal sequestration is only slightly increasing diffusion in cells

Lysosome internal pH of ∼4.5 is maintained by vacuolar proton ATPase (V-H+-ATPase) (de Duve et al., 1974; Kaufmann and Krise, 2007). The low p*K*a of weakly basic small molecule drugs drive their accumulation in the lysosome. There, they become protonated (ionic form is less-permeable), which hinders their back-diffusion to the cytosol. The phenomenon is known as cationic ion trapping (Asokan and Cho, 2002; Kaufmann and Krise, 2007) and results in drug accumulation in lysosomes up to millimolar concentrations (de Duve et al., 1974). We blocked lysosomal acidification using either Bafilomycin A1 or sodium azide prior to drug administration. Bafilomycin A1 is a specific V-ATPase inhibitor and thus inhibits the supply of protons to the lysosomal chamber (Wang et al., 2021). Sodium azide, blocks ATP generation, resulting in the disruption of cellular activities which require ATP (Hiruma et al., 2007; Ishii et al., 2014). As a result, cationic ion trapping is inhibited. These two compounds were used to investigate whether lysosomal trapping is the cause for the 20-40-fold lower diffusion and lower fraction recovery in HeLa cells compared to buffer. Figures S8A and B, and Figure S9A and B show colocalization images for GSK3 inhibitor and quinacrine, respectively (with Lysotracker-633 used as marker) after addition of 100 nM Bafilomycin A1 (Yoshimori et al., 1991). The micrographs show that Bafilomycin A1 strongly reduces lysosomal volumes following quinacrine and GSK3 treatment (Figure S8A *vs* S8B and S9A *vs* S9B). However, D_confocal_ values were not altered much by Bafilomycin A1, as shown in Figures S8C-D and S9C-D for GSK3 inhibitor and Quinacrine, respectively. D_confocal_ for the GSK3 inhibitor is ∼ 0.60±0.02 µm^2^s^-1^, and for Quinacrine ∼ 2.4±0.1 µm^2^s^-1^. This suggests that Lysosomal accumulation of these two drugs is not the reason for the slow diffusion measured in the HeLa cell cytoplasm. It is notable that the D_confocal_ for Quinacrine remained consistent regardless of Bafilomycin treatment, 2 hours after incubation (Fig. S9D, 2.4±0.1 µm^2^s^-1^). However, when measured immediately after injection, the diffusion coefficient was higher at 4.2±0.2 µm^2^s^-1^ (Fig. S5D). This result does not support the notion that the faster diffusion measured immediately after cellular injection relates to lysosomal aggregation, and would better support self-aggregation, or aggregation with other molecules in the cell, which increases over time. This notion is further supported by the almost complete lack in FRAP observed 24 hours after injection (Fig. S5C).

To verify the Bafilomycin results, HeLa cells were pre-treated with Sodium azide at 1 and 50 mM concentrations (Figure 7A and E), followed by treatment with GSK3 inhibitor or Quinacrine (Poste and Papahadjopoulos, 1976). The micrographs in Figures 7A and 7E, show that sodium azide inhibited lysosomal accumulation of these drugs. Comparative FRAP profiles and diffusion coefficients (Figure 7B-D and 7F-H) were slow, but conversely to Bafilomycin, sodium azide treatment did cause a further reduction is rates from D_confocal_ 2.4±0.1 µm^2^s^-1^ to 1.8±0.1µm^2^s^-1^ for quinacrine and from 0.6 to 0.45 µm^2^s^-1^ for the GSK3 inhibitor (Figure 7G and C). Both Bafilomycin and sodium azide treatments resulted in elimination of drug confinement in the lysosome, and the small difference in diffusion rates may be a result of the de-acidification of the lysosomes by sodium azide, which may increase the protons in the cytosol upon treatment.

**Figure 7.**
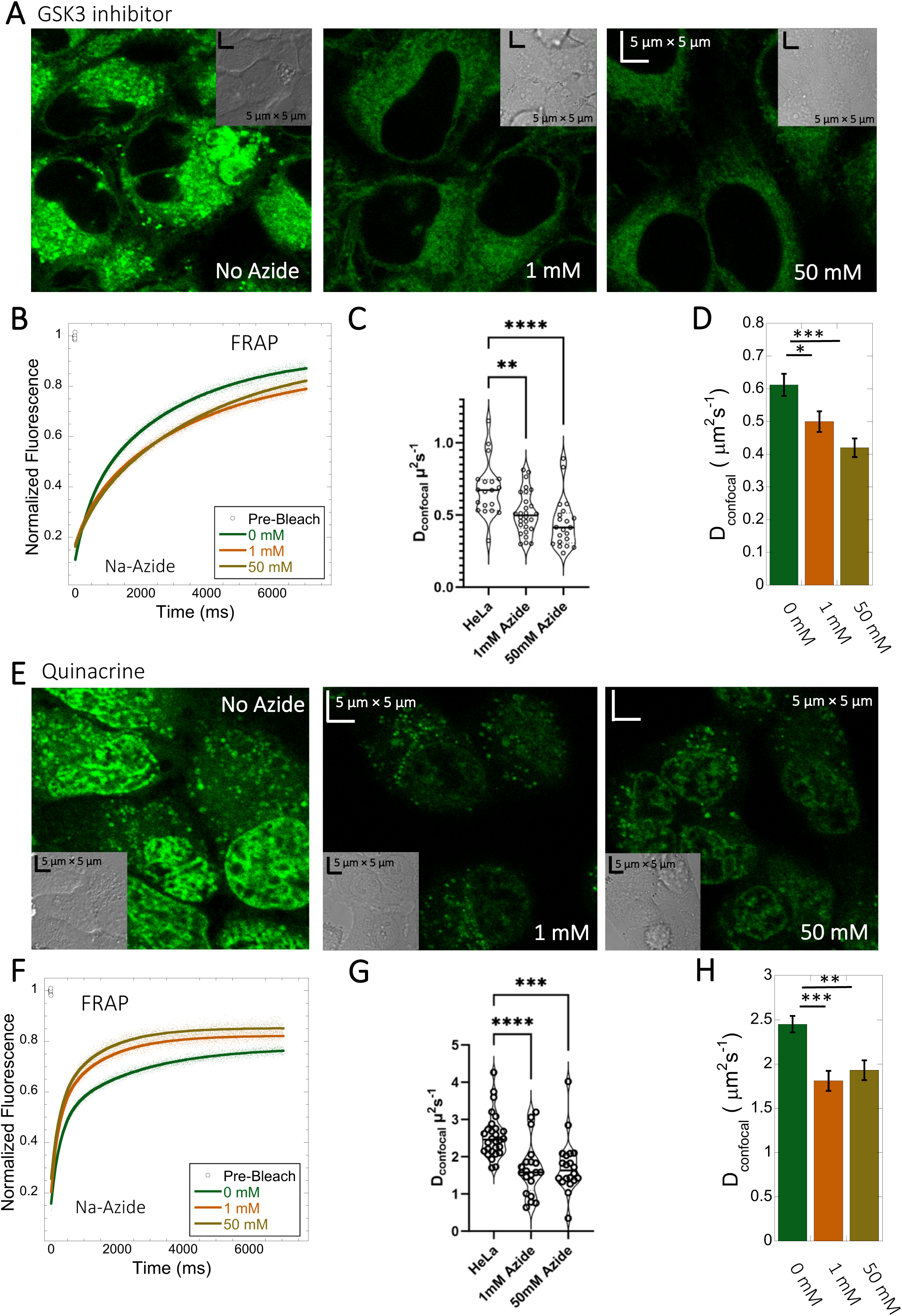
Effect of Sodium Azide on the diffusion of GSK3 inhibitor and Quinacrine inside HeLa cells. (A) GSK3 inhibitor and (E) Quinacrine dihydrochloride treated HeLa cells with or without sodium azide. Comparison of merged FRAP profiles with exponential fits (N= 30; R= 0.99 for each of the fits) and calculated D_confocal_ values from individual fits and combined averaged fit for (B-D) GSK3 inhibitor and (F-H) Quinacrine dihydrochloride are shown. D_confocal_ estimations from individual FRAP curves are shown in (C,G). Error bars represent SE calculated from fitting the FRAP progression curves, which are averaged over at least 30 independent measurements. Statistical significance calculations are detailed in the materials and methods section.

### 5-amino fluorescein diffuses slowly in HeLa cells and is accumulated within lysosomes

To experimentally test the relation between the p*K*a of small molecules and their in-cell diffusion, we measured the diffusion of 5-amino fluorescein (AM-Fluorescein, M.W.= 347.3 Da, Figure 8A), which is structurally similar to fluorescein disodium salt, but with an additional -NH_2_ group that increases its p*K*a from ∼3 to ∼9 (Figure 8A). Figures 8B and C show comparative FRAP traces for these molecules, and for CCF2. While fluorescein disodium salt has a fast FRAP rate, and is completely recovering, the cationic charged AM-fluorescein recovers much slower with a fractional recovery of 0.4. Moreover, while fluorescein disodium salt is not sequestered in lysosomes, 5-Amino fluorescein is. This sequestration is alleviated by adding sodium azide to the cells (Figure 8D) but has only a small effect on the FRAP rate and fractional recovery (Figure 8B). This experiment strongly suggests that the functional primary amine group is the main contributor towards the slow diffusion and low FRAP recovery of AM-fluorescein in the cell cytoplasm.

**Figure 8.**
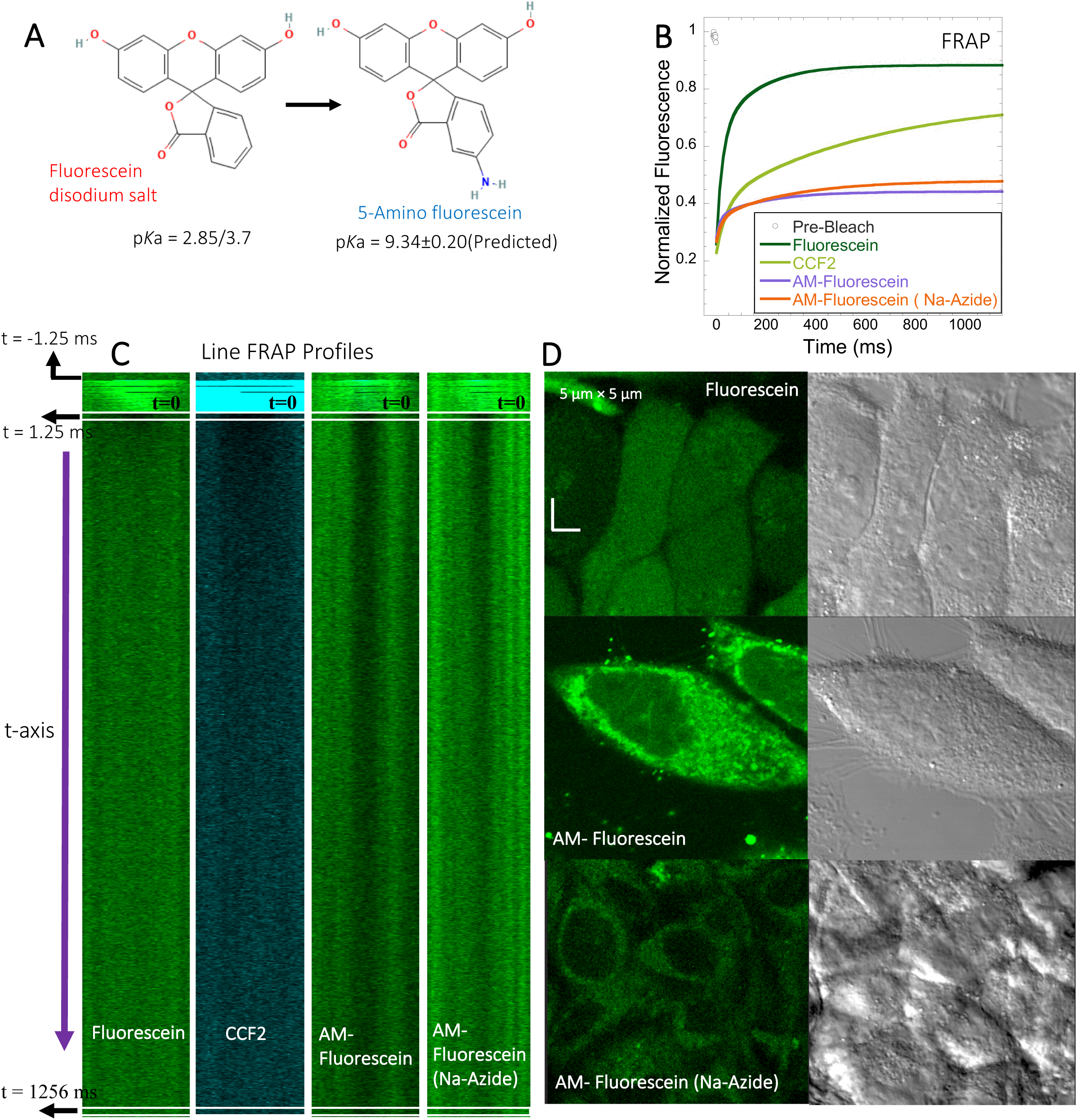
FRAP of Fluorescein analogues inside HeLa cells. (A) Structures of Fluorescein analogues with p*K*a values. (B) FRAP recoveries with exponential fits for Fluorescein, CCF2 and 5-amino Fluorescein, with/without Na-Azide treatments in HeLa cells (N= 30; R= 0.99 for each of the fits) . (C) Line FRAP profiles with time lapses. Micrograph images of treated HeLa cells after small molecule incubations.

### Acidification of primaquine and BODIPY increases their diffusion and FRAP recovery

We substituted the NH_2_ of primaquine (antimalarial drug) and BODIPY compounds for the charge-neutral-NHAc functional group (Figure S10A and S11A). Due to their low quantum yield, we could not implement the faster Line-FRAP protocol, therefore, we used standard FRAP, which suits the slow diffusion of both compounds. As seen from the FRAP profiles (time-dependent monographs) in Figures S10B-D for primaquine derivatives and in Figures S11B-C for BODIPY derivatives, the recovery is much slower for free amine-containing moieties. Figure S10I-J shows that the fractional recovery for primaquine is only ∼0.27, whereas, for primaquine-NHAc, the recovery is ∼0.6. Comparing the measurements using micro-injecting versus incubation of the cells for FRAP measurements for primaquine-NHAc gave similar results (Figure S10E-H). Unfortunately, the classical FRAP is not fast enough to calculate the diffusion coefficients, as the dead time was 150 ms. For the BODIPY analogues, the observations were similar, but to a smaller extend. For BODIPY-NH_2_, the recovery percentage is ∼57%, whereas, for BODIPY-NHAc, the recovery is ∼67% (Figure S11H). The estimated recovery half-life also supports the faster diffusion of BODIPY-NHAc, as shown in Figure S11I.

### Basic small molecules do not colocalize with lipid droplets, the ER or nucleic acids

To identify the location of colocalization of basic small molecules, we tested colocalization of GSK3 inhibitor in lipid droplets using the Nile red dye (Figure S12) and in the ER using a specific BFP/mCherry tagged fluorescent antibody markers (Figure S13). Next, we looked for colocalization of Mitoxantrone with nucleic acids present in cytoplasm using SYTO blue (Figure S14). No colocalization was found between the drugs and either lipid droplets, the ER or nucleic acids.

Super resolution images of GSK3 inhibitor treated HeLa cells were taken to improve the spatial distribution (Figure S15); which however, this did not contribute new information to the puzzle. Therefore, we could not pinpoint the location of sequestration of weakly basic small molecule drugs within the cellular cytoplasm, except the lysosome.

## DISCUSSION

In this study, we selected well-known and widely used small molecule drugs to study their behavior in aqueous solutions and living cells. All drug molecules used here are fluorescent, allowing for their FRAP measurements. Previously, we found that small-molecule drugs diffuse differently in crowded media than in a simple buffer solution, even if aggregation is not an issue (Dey et al., 2022). Here, we show that many small molecules get trapped within the cellular cytoplasm, resulting in extremely slow diffusion and lower fractional recovery after FRAP. This is particularly observed for cationic-charged small molecules, with D_confocal_ values of 20-40-fold lower than observed for fluorescein or CF514 (Figure 9A vs 9B), which, in many cases is accompanied by low recovery after photobleaching. The later suggests that most molecules are occluded to components of the HeLa cell cytoplasm. Even dense cell extract from HeLa cells is not mimicking the *in vivo* results (Figure 4A-C and 5A-C). This is very different from protein diffusion, where none of the 16 proteins measured here showed slow diffusion or low fractional recovery, despite their *E.coli* origin, expressed in HeLa cells. Slow diffusion and lower fractional recovery were limited to basic small molecules, showing D_confocal_ values of 0.2-2 µm^2^s^-1^. While the fractional recovery for all proteins is ∼0.9, for many basic small molecules it is 0.2-0.5.

**Figure 9.**
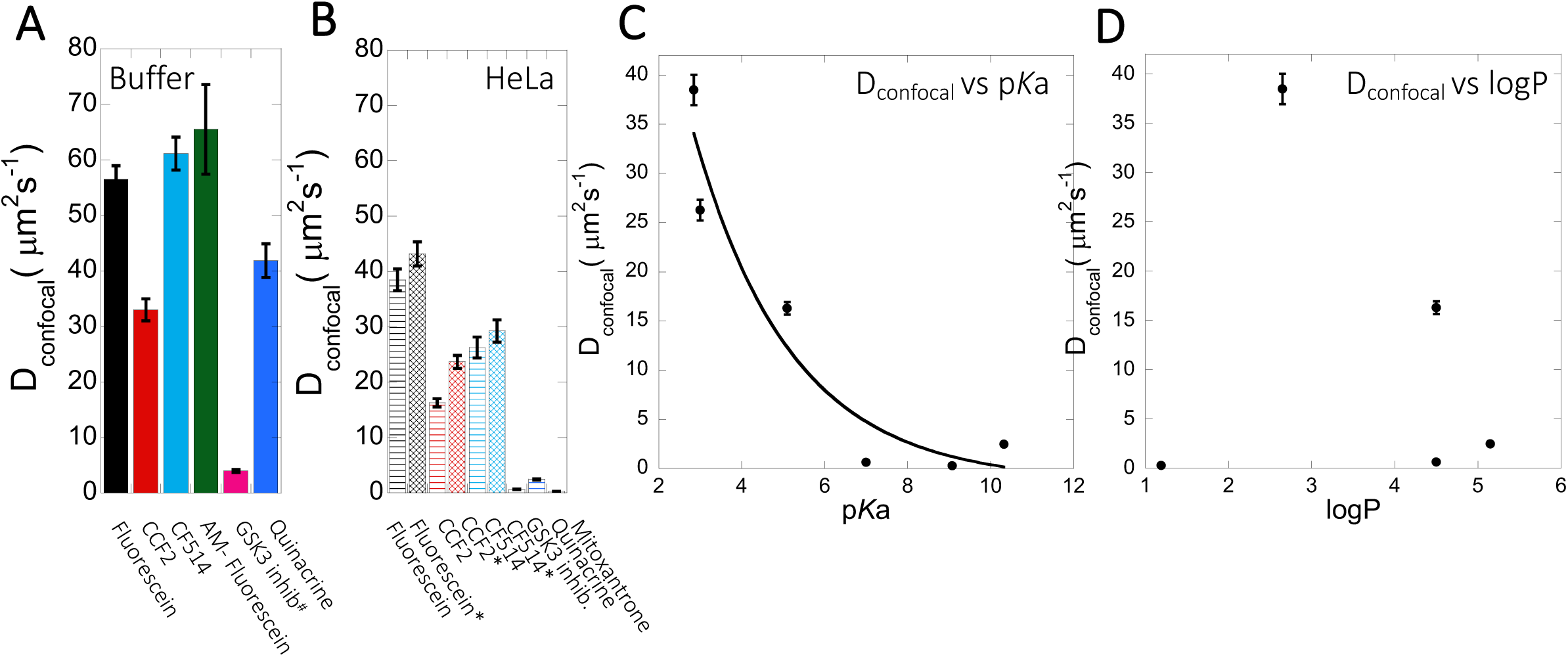
D_confocal_ of small molecules in PBS buffer and inside HeLa cells. Comparison of D_confocal_ values of small molecule drugs in (A) PBS buffer and in (B) HeLa cells. Error bars represent SE calculated from fitting the FRAP progression curves, which are averaged over at least 30 independent measurements. Dependency of D_confocal_ values on (C) p*K*a and (D) logP values in HeLa cells. ^#^ in (A) denotes DMEM media (instead of PBS), * in (B) denotes measurements done in the nucleus.

As primary culprit for the slow diffusion of the basic small molecule drugs we suspected sequestration in the lysosomes. Indeed, high level of accumulation in the lysosomes were found for all the basic drugs. However, pre-treatment with sodium azide or Bafilomycin A1, which we showed to inhibit lysosomal accumulation, had no or only a small effect on diffusion or fractional recovery after FRAP, which stayed low. Therefore, we evaluated colocalization of GSK3 inhibitor with lipid droplets, the ER or nucleic acids present in cytoplasm (Figure S12-14), but did not detect any.

The conclusive evidence that slow diffusion and lower FRAP recovery results from the drugs being basic came by altering three molecules from being basic to acidic and vice versa. The best example here is fluorescein, where transformation to 5-amino fluorescein drastically reduced its diffusion as well as its fractional recovery after FRAP. Two additional examples here were Primaquine that was modified to Primaquine Acetate and BODIPY-NH_2_ that was modified to BODIPY-NHAc. In both cases, the diffusion became faster, and the fractional recovery increased significantly. We did not find any specific correlation between individual average diffusion rate constants with the concentration of small molecule drugs inside the cell (Figure S3A-D). For fraction recovery, we saw some decrease at high concentrations for the GSK3 inhibitor and quinacrine, but not fluorescein (Figure S3E-H), which would support out notion that they are sequestered in the cell. Correlating the calculated D_confocal_ values with p*K*a shows a clear trend, where diffusion is strongly slowed down at higher p*K*a values (Figure 9C). Conversely, logP, molecular weights or number of aromatic rings present in the drug molecules did not relate to D_confocal_ values (Figure 9D and S16A-B). This finding is important, as the activity of a drug within the cell is dictated by its active concentration. When the drug is sequestered, its active concentration is reduced. Even, if it is not fully sequestered, but only diffuses much slower, it will result in lower association rate constant, and thus lower affinity towards the drug target. As a result, to keep the drug active, its dose has to be increased, which can have negative implications on side effects, due to off-target binding.

## CONCLUSIONS

Our study shows that *in vitro* biophysical crowding studies for small molecule drugs are of limited value if one wants to understand the biophysical behavior of the same drug within the cell. While it is true that availability of a drug within a cell is only one factor dictating its biological activity, it is a crucial one. Our findings here raise an important limitation on the standard rules for drug design, as these do not consider the stickiness of basic small molecules within the cell. We were able to directly address this question by using fluorescent molecule drugs, and measuring their location and diffusion within cells. Most importantly, we also show that by blocking protonation of a number of basic compounds we were able to increase substantially their diffusion and recovery after photobleaching, and thus potentially their activity. These findings may be consequential in future drug development.

## MATERIALS AND METHODS

All the reagents used are described in a table format (Table S2, Key resources). The supplementary method sections describe the synthetic procedures, characterizations, and purity of the synthesized compounds.

### Selection of drugs used in this study

To identify small molecule drugs that have fluorescence in the visible light, we screened in high-throughput 384 well plates 1600 commonly used, small molecule drug compounds for their penetration into HeLa cells and fluorescence at the DAPI (377/447), FITC (475/520), TexasRed (560/624) and Cy5 (631/692) channels. As positive control, we used Pyrvinium (red), Doxorubicin (green)and CSB (blue). Out of those, 16 compounds gave positive results. These compounds were then verified for their photostability, availability and solubility. This resulted in the list of compounds shown in Table 1, which includes also compounds used by us in previous studies.

Mammalian Cell Culture. HeLa cells were grown in 35-mm glass-bottomed dishes (MatTek Corporation) in DMEM (1X) Gibco (Life Technologies Limited) supplemented with 1X pyruvate, penicillin/streptomycin (BioIndustries), and 10% fetal bovine serum (Life Technologies Limited). The cells were subcultured when 80% confluence was reached using trypsin-EDTA for cell detachment. 2x10^5^ HeLa cells in 2.5 ml of DMEM were pipetted into glass-bottomed dishes and incubated overnight. The cells were cultured in a humid atmosphere at 37°C and 5% CO2. The cells were imaged 24–30 h after seeding. Before the microinjection, the medium was aspirated, and fresh medium was supplemented with 25 mM HEPES, pH 7.4. For drug treatment, HeLa cells were incubated with 10 µM of drug (diluted 1:1000 from stock) for indicated times at 37 °C, followed by three times PBS 1X wash prior to imaging. For dose-dependent studies, the drug concentrations varied typically between 2-12 µM. Sodium azide concentration was 1 and 50 µM. Bafilomycin A1 was used at 100 nM concentration. Similarly, during the colocalization study, HeLa cells were treated with LysoTracker dye (1X) with 1:1000 dilution for 30 mins to 1 h at 37 °C, followed by 3x washing with PBS. Quinacrine, GSK3 inhibitor, Mitoxantrone, 5-amino Fluorescein, Amidoquine, Primaquine and BODIPY analogues were incubated with cells, while Fluorescein, Quinacrine, Primaquine-Ac, and CF514 dye were also micro-injected inside the HeLa cells.

### Microinjection into HeLa cells

Microinjections were performed using the Eppendorf FemtoJet microinjector attached to the Eppendorf InjectMan NI2 micromanipulator. The fluorescein sodium salt, CCF2, CF514, quinacrine, primaquine analogues, BODIPY analogues were dissolved in DMSO and highly concentrated stock aliquots were made. Diluted PBS solutions of small molecules were injected into cells using glass capillaries from Warner instruments and pulled by a vertical puller (Narishige). For every measurement, a single pressure pulse was applied to deliver the sample into the cell. Air was administrated at 15– 25 hPa for 0.1-0.3 s. For injections, single cells containing morphologically healthy and well-connected HeLa cells were selected. Before and after the microinjection, cell morphology and membrane integrity were confirmed by visually inspecting the injected cells.

### HeLa cell extract preparation

Cytoplasmic HeLa cell extracts were prepared as described previously with slight modifications (Zotter et al., 2017). HeLa cells from 4×10 cm plates at 80% confluency were washed, treated with trypsin and collected. Pellets were mixed with 500µl of RIPA buffer, IMP40, 5 µL of protease inhibitor and stored in ice. After 20 min, the mixture was centrifuged at 13k×g. This process was repeated twice to obtain concentrated extract. A BSA calibration curve determined the total protein concentration; the final concentration was up to 100 mg/mL. After liquid nitrogen freezing, the final cell extract solution was stored at -80° C.

### Protein purification and dye labeling

Bacterial proteins used in this study were purified as described by us previously (Marciano et al., 2022). The dye labelling procedure with CF514 for purified proteins and removal of excess dye was described in detail (Dey et al., 2021).

### Confocal microscopy and FRAP Analysis

*Confocal microscopy.* Images were collected with an Olympus IX81 FluoView FV1000 Spectral/SIM Scanner confocal laser-scanning microscope, using 60X DIC oil-immersion objective, N.A. 1.35. For fluorescein sodium salt fluorescence measurements, excitation was done at 440 nm, using a diode laser at an output power of 1-4 % of maximal intensity for high to low concentrations. In contrast, emission was recorded from 520 to 550 nm using the spectral detection system. For CCF2-FA, excitation was done with the laser at 440 nm using 1-2 % of the maximal intensity, while emission was collected from 470-560 nm with SDM560 emission dichromator as cut-off filter. For quinacrine DHC and GSK3 inhibitor, excitation was done at 488 nm laser using 1-2% of the maximal intensity, while emission was collected from 502-560 nm with SDM560 emission dichromator as cut-off filter. During colocalization experiments of quinacrine DHC and GSK3 inhibitor with LysoTracker-633 dye, the same setting was used for detecting the small molecule drugs in Channel 1 (green). LysoTracker-633 dye was detected in Channel 2 (red) by exciting at 635 nm laser using 1-2 % of the maximal intensity, while emission was collected from 655-755 nm. Colocalization for Mitoxantrone (red channel) was done by using quinacrine to track the Lysosomes (green channel). Percentage of volume and drug intensity (material) above threshold is defined as percent colocalized (Figure S16C-D). For mitoxantrone and BODIPY analogues, excitation was done at 635 nm laser using 1-2 % of the maximal power, and emission was collected from 655-755 nm in Channel 2 (red). In the blue channel, excitation was done by 400 nm laser using 2-4% of maximal power, whereas the emission was collected from 420-480 nm with SDM480 emission dichromator as cut-off filter. The same setting is used in the blue channel for the SYTO blue maker (to visualize free nucleic acids) and blue fluorescent protein marker (BFP to visualize the ER). For the mcherry protein marker (to visualize the ER), the same setting for the red channel (Channel 2) is used. For every colocalization experiments, proper blank experiments were performed separately in all the respective channels with the same microscope settings to avoid any bleeding/leakage of emission signals from one Channel to another Channel, as shown in Figure S17. Image analyses were performed using FluoView/Imaris software, and data analyses were performed using Kaleidagraph software version 4.1 (Synergy). Colocalization coefficients as a percentage of material/volume above threshold colocalized are calculated using Imaris software version 7.

### Line-FRAP and Classical XY-FRAP

Line-FRAP was carried out in liquid drops and HeLa cells. For photobleaching, “Tornado” of 4 pixels (2x2, 1-pixel = 0.207 micrometer) as default laser settings by the Olympus Fluoview confocal system (Dey et al., 2021). It is the smallest area achievable using Tornado. The bleach circle was kept precisely in the middle of the scanning line. The lines we scanned were unidirectional with time intervals of 1.256 ms 1000 times (equivalent to 1.256 s) in the majority of the measurements. The number of scans prior to, during, and after the photobleaching was 10, 42, and 948, respectively. Only in some specific cellular measurements (for GSK3 inhibitor and Quinacrine DHC, where recovery is very slow) the unidirectional lines were scanned 5000 times. Photobleaching was achieved by using the laser at 405-nm or with 635-nm excitations for the 63-millisecond duration at full intensity (100%). The simultaneous scanner moved at a 100 µs/pixel speed to perform an efficient photo-bleach. We have used two simultaneous scanners during the Line-FRAP experiments: one scanner (at 405 nm with the full intensity of 100%) for photobleaching and another scanner (at 440/515/635 nm with weak intensity) for data acquisition. For all the drugs except Mitoxantrone and BODIPY analogues (emission in red wavelengths), photo-bleach was performed by 405 nm laser. For fluorescence signal detection: fluorescein 440 nm laser (1-4%); CCF2 440 nm laser (1-2%); GSK3 inhibitor 440 nm laser (0-2%); quinacrine (1-2%) of maximal intensity were used. Emission collections were done from 520-550 nm for quinacrine DHC and GSK3 inhibitor.

Line FRAP was not possible for primaquine analogues, as it suffers from strong photo-bleach during recovery. Here classical XY FRAP was performed, with photo-bleach done by 405 nm laser with full intensity. The frame rate is maintained at 172 ms with a bleach pulse duration was 150 ms. Due to microscope setting limitations, Line FRAP is also not feasible for Mitoxantrone and BODIPY analogues. A classical XY-FRAP using two simultaneous scanners is done for these molecules. Here, bleach was performed by simultaneous 635 nm laser with the full intensity of 100%, and the main excitation was done by 635 nm laser (with the power of 4%). Emission collections were done from 655-755 nm. For BODIPY analogues FRAP recoveries, the bleach pulse time was 150 ms. The bleach pulse time was also 150 ms for primaquine analogues for comparative FRAP profile collections. The frame rate is maintained at 172 ms. Using the Olympus IX81 FluoView FV1000 Spectral/SIM Scanner confocal laser-scanning microscope, using Tornado (which requires SIM scanner to be loaded) greatly enhances bleaching efficiency. In addition, it shortens the time to obtain the first measurement after bleach (which is immediate in this mode). This property is highly beneficial for Line-FRAP measurements, where the time scale of data acquisition plays an important role. The Fluoview SIM scanner unit synchronizes laser light simulation with confocal and multiphoton imaging to avoid interruption to image observation during laser stimulation or manipulation. We have varied the intensity of the lasers to achieve an optimal signal/noise ratio. Fluorescence recovery plots were fitted to a double exponent growth curve. FRAP experiments were also performed inside the PBS buffer drops and crowding conditions. Line FRAP was done on the cells taken from different plates, treated independently. A coverslip was introduced at the top of the drop to stop evaporation during measurements. Our previously developed method on Line FRAP protocol has been employed to calculate diffusion coefficients from the FRAP rates and averaged bleach sizes. Diffusion rates cannot be calculated using XY-FRAP. However, FRAP parameters were kept constant so that qualitatively we could compare the average half-life and percentage of recoveries. When the areas of bleach were selected in the drug-treated cell cytoplasm, we avoided the lysosomes as much as possible, within the resolution limits of the confocal microscope. Lysosomes themselves were measured to move within the cytoplasm with an diffusion coefficient of 0.03-0.071 *µ*m^2^ s^−1^ (Bandyopadhyay et al., 2014), which is much slower than the diffusion measured for even the slowest compounds using fast Line FRAP, further validating that we did not measure lysosome diffusion.

### Determination of D_confocal_ from FRAP results

We previously developed (Dey et al., 2021) the FRAP in Line mode (Line FRAP) method to monitor the diffusion rates of proteins in various environments. Line-FRAP allows a much faster data acquisition rate compared to classical XY FRAP. It is more suitable for measuring diffusion rates for fast diffusing molecules. To calculate D_confocal_, 20-30 independent measurements on different cells (n) were binned, and the curve fit of the progression curve was used to obtain t_1/2_, and r_e_ values and their associated errors as determined from the fit of the curve. r_n_ was obtained from bleaching a fixed sample. The D_confocal_ values were then calculated using equation 1 (see introduction and (Dey et al., 2021)). The standard errors (SE) of the individual parameters were combined to obtain D_confocal_±SE. The errors and means were used to calculate the significance level of the difference between two treatments, with ns for non-significant, * being for 0.05-0.01, ** being for 0.01-0.005, *** for 0.005-0.001 and **** <0.0001. D_confocal_ values from individual Line FRAP profiles were calculated the same way, and one-way ANOVA statistical tests were performed to calculate the significance between treatments using the standard function available in GraphPad Prism v10.1 using Dunnett for multiple comparisons. To verify SE values, we repeated the ∼20-30 measurements independently multiple times, which gave the same SE as obtained from individual curve fits of n = 30 cells.

## ASSOCIATED CONTENT

### Supporting Information

The Supporting Information is available free of charge on the eLife website.

## AUTHORS

Debabrata Dey, Department of Biomolecular Sciences, Weizmann Institute of Science, Israel.

Shir Marciano, Department of Biomolecular Sciences, Weizmann Institute of Science, Israel.

Anna Poryvai, Institute of Organic Chemistry and Biochemistry of the Czech Academy of Sciences, Flemingovo náměstí 542/2, Prague 6, 160 00 Czech Republic

Ondrej Groborz, Institute of Organic Chemistry and Biochemistry of the Czech Academy of Sciences, Flemingovo náměstí 542/2, Prague 6, 160 00 Czech Republic

Lucie Wohlrábová, Institute of Organic Chemistry and Biochemistry of the Czech Academy of Sciences, Flemingovo náměstí 542/2, Prague 6, 160 00 Czech Republic

Tomás Slanina, Institute of Organic Chemistry and Biochemistry of the Czech Academy of Sciences, Flemingovo náměstí 542/2, Prague 6, 160 00 Czech Republic

### Author Contributions

D.D. performed all the experiments. S.M. purified all the bacterial proteins. A.P, O.G., L.W., and T.S. designed the synthetic targets and synthesized primaquine acetate and BODIPY derivatives. D.D. and G.S. conceptualized the idea, analyzed the data and wrote the original draft of the manuscript. D.D., G.S., and T.S. reviewed and edited the paper. G.S. supervised the project. The manuscript was written through contributions of all authors. All authors have given approval to the final version of the manuscript.

### Funding Sources

The Israel Science Foundation grant number 1268/18 (GS). The Czech Science Foundation (project No. 22-20319S) for funding this project (T.S.)

## ACKNOWLEDGMENT

We would like to thank Dr. Ori Avinoam from Department of Biomolecular Sciences, and Prof. Hagen Hofmann from Department of chemical & structural Biology, Weizmann Institute of Science for their valuable discussions and advices. We want to acknowledge Dr. Reinat Nevo for her help with Imaris software to analyze the microscopy images.

## ABBREVIATIONS

FRAP: fluorescence recovery after photo-bleaching
CCF2: coumarin cephalosporin fluorescein
BODIPY: boron dipyrromethene analogues
NMR: Nuclear magnetic resonance.

---

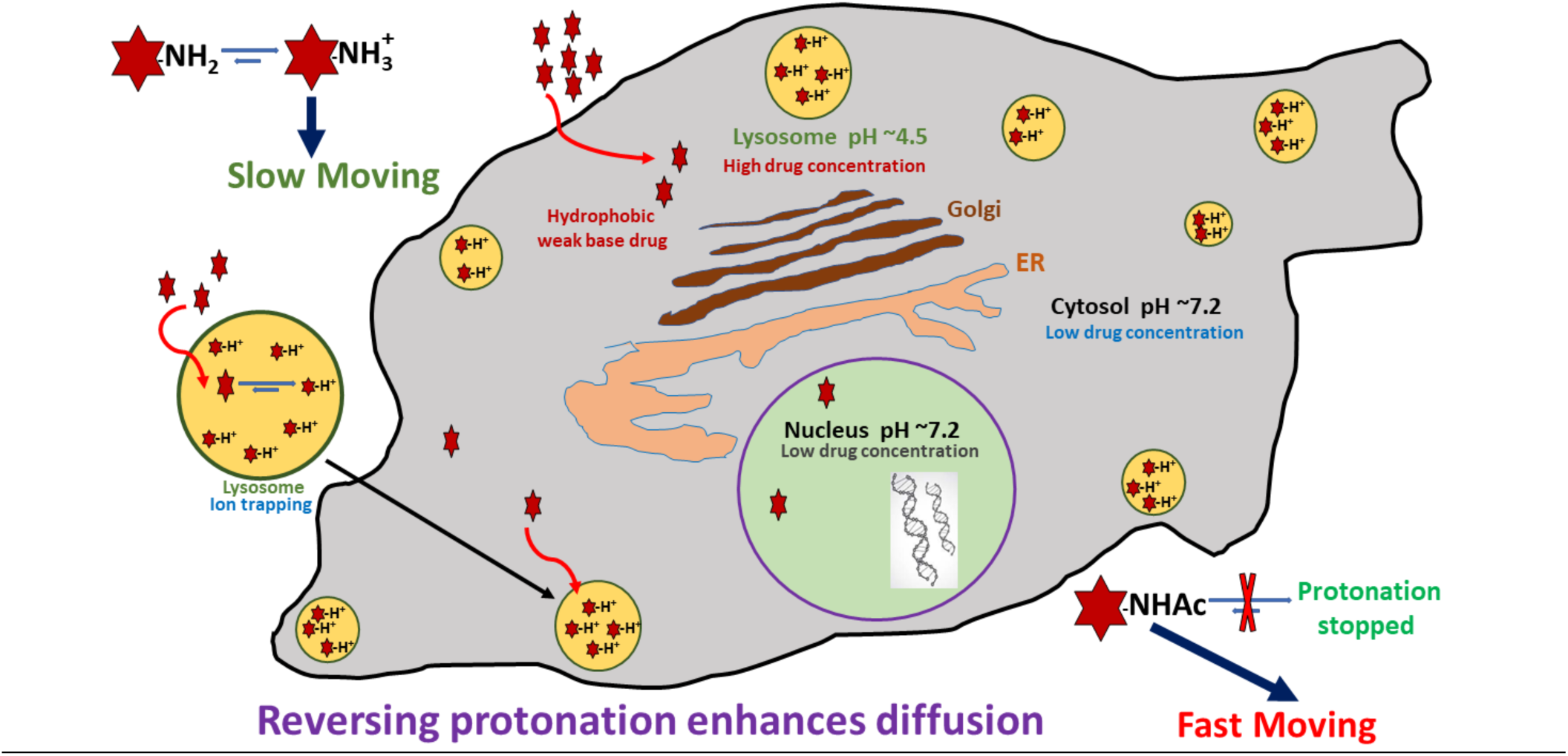

## Supplementary Information

**Table S1:** Oligomeric state, Isoelectric point and Net charge at pH 7.4.

| Protein entry | Isoelectric point: pI | Net charge at pH 7.4:<br>*Z* | *D*<sub>confocal</sub> | Oligomeric state | Uniprot (kDa)<br>Monomer |
| --- | --- | --- | --- | --- | --- |
| NfuA | 4.534 | -15.767 | 24.8±1.1 | M/D | 20.93 |
| NadK | 6.387 | -4.808 | 29.8±1.4 | M/D/T | 32.57 |
| FabG | 6.250 | -1.66 | 30.33±1.4 | D/T/H | 25.56 |
| E-fts | 5.228 | -9.527 | 22.5±1.0 | M/D | 30.29 |
| SodA | 6.511 | -2.481 | 25.5±1.2 | D | 22.97 |
| tpiA | 5.688 | -7.851 | 27.9±1.7 |  |  |
| GpmA | 5.913 | -4.262 | 21.7±1.0 | D | 28.43 |
| Can | 6.230 | -5.908 | 18.7±1.4 | T | 25.1 |
| Upp | 5.335 | -6.829 | 23.0±1.1 | M/D | 22.53 |
| SpeB | 5.160 | -14.788 | 30.4±1.4 | Tri/H | 33.56 |
| IspD | 6.257 | -5.445 | 24.1±1.2 | D/T/H | 25.61 |
| BaeR | 5.554 | -6.032 | 21.9±1.1 | M/D | 27.66 |
| Acul | 5.669 | -5.986 | 28.0±1.3 | M/D/T/H | 34.73 |
| PyrF | 5.874 | -3.9 | 26.0±1.20 | M/D | 26.35 |
| ThiD | 5.793 | -6.842 | 28.7±1.3 | M/D | 28.64 |
| NadE | 5.603 | -4.993 | 23.2±1.1 | D | 27.16 |
| CRP | 7.979 | +1.388 | 30.1±1.3 | D/T | 23.64 |
| DES3im-4rp4 | 5.041 | -5.659 | 29.4±1.4 |  |  |
| DES7im-3rk6 | 4.599 | -10.654 | 28.8±1.3 |  |  |
| 6E5C_denovobeta30 | 6.336 | -0.714 | 30.0±1.4 |  |  |
| BSA | 4.5–4.8 | -15 | 29.6±1.4 | M/D | 66.5 |

Isoelectric points and net-charges were predicted using Prot Pi. Oligomeric states were taken from (Marciano et al., 2022) native mass spectrometry measurements.

**Table S2.** Key resources table.

| REAGENT or RESOURCE | SOURCE | IDENTIFIER |
| --- | --- | --- |
| Chemicals, peptides, and recombinant proteins |  |  |
| Dulbecco's phosphate-buffered saline (PBS 1X) | Biological Industries | Cat# 02-023-1A |
| HEPES buffer | Fisher BioReagents | LOT# 170358 |
| DMEM(1X) Gibco | Life Technologies Limited | REF# 41965-039 |
| Fetal Bovine Serum (Gibco) | Life Technologies Limited | REF# 12657-029 |
| Trypsin-EDTA Solution A | Biological Industries | REF# 03-050-1B |
| Penicillin/ Streptomycin | Biological Industries | REF# 03-031-1B |
| Sodium Pyruvate Solution | Biological Industries | REF# 03-042-1B |
| 35-mm glass-bottomed dishes | MatTek Corporation | P35G-0-14-C |
| Pierce™ Dye Removal Columns | Thermo-Fisher scientific | Cat# 22858 |
| Gebaflex tubes 3.5KDa (GeBa) | TIVAN BIOTECH | Cat# MIDI3-100 |
| Capillary Glass Tubing | Warner Instruments | Model No. G120TF-4<br>203-776-0664 |
| 96- well Black F-Bottom Plate | Greiner-bio-one | REF# 655076 |
| Disposable cuvettes | Fisher Scientific | ZEN0040 |
| BSA (Albumin Bovine, fraction V) | MP Biomedicals, LLC | CAS# 9048-46-8 |
| HEWL (Lysozyme from chicken egg white) | Merck | CAS# 12650-88-3 |
| Myoglobin (from equine heart) | Merck | CAS# 100684-32-0 |
| Doxorubicin | AdooQ Bioscience | Cat# A14403 |
| Fluorescein disodium salt | chemcruz | Cat# sc-206026 |
| GSK3 inhibitor SB216763 | Abcam | Cat# ab120202 |
| Quinacrine dihydrochloride | Abcam | Cat# ab120749 |
| Primaquine biphosphate | Merck | CAS# 63-45-6 |
| CCF2-FA | Thermo-Fisher Scientific | Cat# K1039a |
| CF514 Labelling dye | Biotium | CF514 Dye<br>Alternative green fluorescent dye |
| Amodiaquine | AdooQ Bioscience | Cat# A17660 |
| Sodium Azide | Merck | CAS# 26628-22-8 |
| Mitoxantrone | Abcam | Cat# ab141041 |
| Bafilomycin A1 | Abcam | Cat# ab120497 |
| 5-amino Fluorescein | Abcam | Cat#ab145305 |

### Characterization of Prmaquine (Prim) and N-acetylated Primaquine (PrimAc)

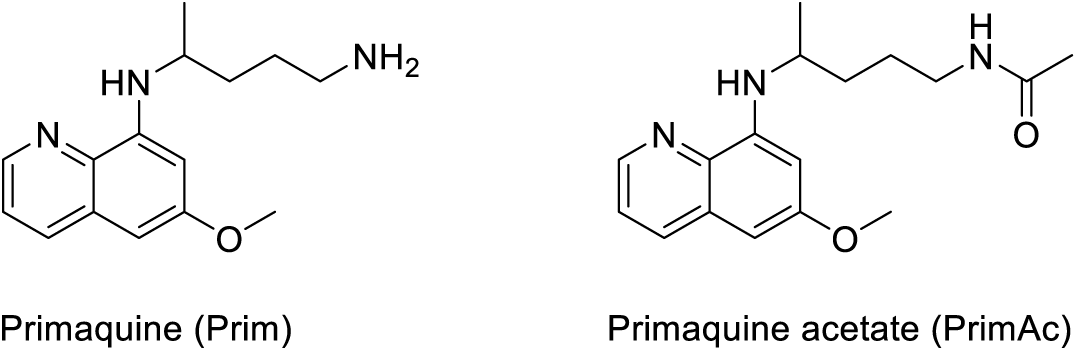

The absorption and emission spectra of Primaquine analogues are shown in Figure S18 A& B respectively. They were measured in the mixture of 10% DMSO in PBS or in Glycine/NaOH buffer.

#### Materials & Reagent

Glycine/NaOH (pH = 10) (preparation: 80 mL of distilled water, 0.6 g of glycine, 0.25 g of NaOH, pH adjusted to 10 using 35% HCl, diluted to 100 mL)

#### Synthesis of PrimAc

To the stirred solution of Primaquine bisphosphate (100 mg, 220 µmol, 1 eq) and TEA (92 µL, 660 µmol, 3 eq) in DMF acetic anhydride (21 µL, 220 µmol, 1 eq) was added at RT. Reaction mixture was stirred for 5 minutes and the reaction mixture was diluted with dichloromethane (50 mL) and extracted 2x with water. Water phase was extracted with 20 mL of DCM and combined organic layers were extracted with brine. The organic phase was evaporated to the volume of approximately 1 mL and the solution was liquid loaded to the column. Gradient chromatography was done using cyclohexane, ethyl acetate and methanol (grad.: 100% cyclohexane – 30% cyclohexane, 60% ethyl acetate, 10% methanol). The chromatography was repeated to obtain the target compound as yellowish oil in 94% yield and 98.5% purity.

**^1^H NMR** (401 MHz, CDCl3) δ 8.53 (dd, *J* = 4.2, 1.6 Hz, 1H), 7.92 (dd, *J* = 8.3, 1.6 Hz, 1H), 7.31 (dd, *J* = 8.2, 4.2 Hz, 1H), 6.34 (d, *J* = 2.5 Hz, 1H), 6.28 (dd, *J* = 2.6, 0.6 Hz, 1H), 5.99 (d, *J* = 8.4 Hz, 1H), 5.50 (s, 1H), 3.69 – 3.58 (m, 1H), 3.36 – 3.18 (m, 2H), 1.92 (s, 3H), 1.72 – 1.63 (m, 5H), 1.30 (d, *J* = 6.4 Hz, 3H). **^13^C NMR** (101 MHz, CDCl3) δ 170.03, 159.45, 144.92, 144.37, 135.36, 134.84, 121.91, 96.84, 91.75, 55.24, 47.85, 39.61, 34.01, 26.30, 23.33, 20.65. (Figure S19 A&B). HRMS spectrum of Primaquine-Ac is shown in Figure S20.

#### Hydrolytic experiment

The compound PrimAc (1 mg) was dissolved in 20 µL DMSO and diluted with 1.98 ml 0.1M PBS. The solution was measured by HPLC-MS (0-day, 1 day, 7 days, 1 month). The sample was stored in the dark at 25 °C between measurements. (Figure S18 C&D)

### Characterization of BODIPY-NH2 (BDP-NH2) and BODIPY-NHAc (BDP-NHAc)

The Structures and photo-physical properties of BODIPY analogues are shown in Figure S21A. The absorption and emission spectra of BODIPY-NH2 and BODIPY-NHAc in PBS/DMSO are shown in Figures S21 B&C, respectively.

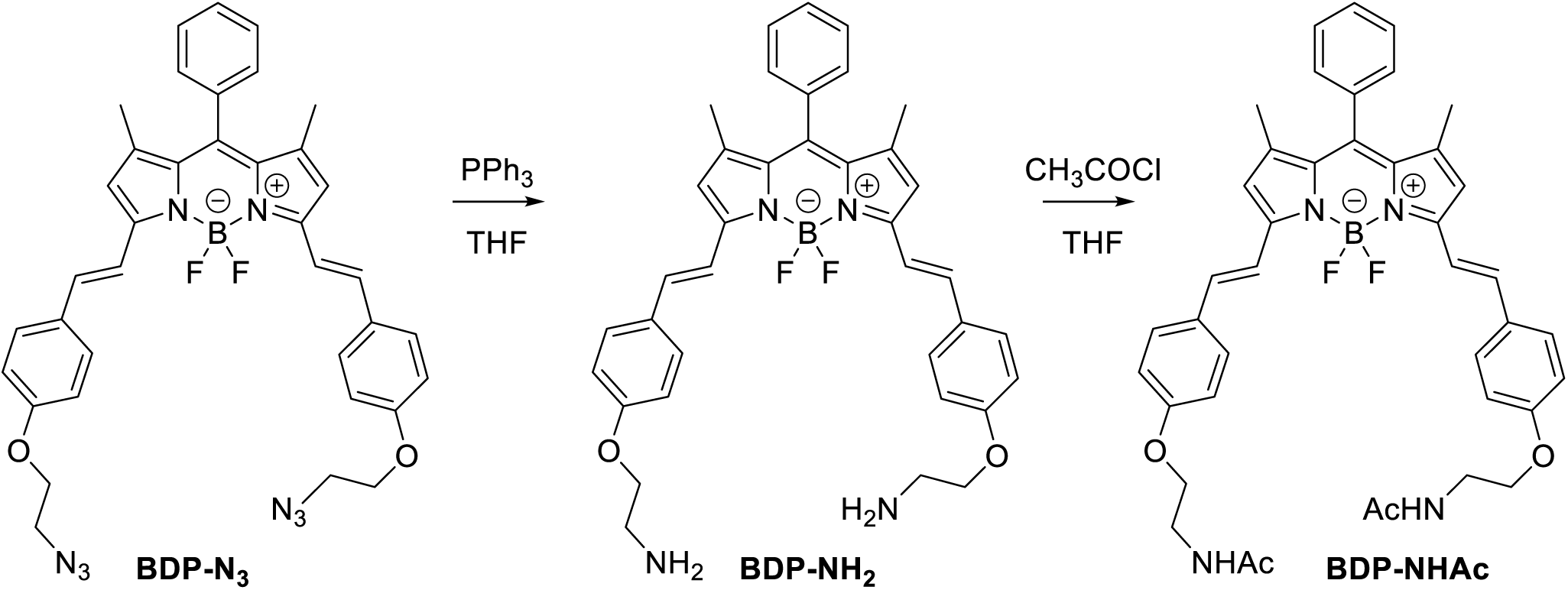

**BDP-NH2**: To a solution of a starting **BDP-N3** (60 mg, 0.1 mmol, 1 eq.) and PPh3 (47 mg, 0.2 mmol, 2 eq.) in THF (8 mL) water (1 mL) was added. The reaction mixture was stirred for 24 h and transferred on 20 g of silica. First fractions were eluted using a MeOH/DCM, 6/1 mixture followed by elution of the product using MeOH/DCM/NH3 aq, 90/7/3, mixture. Organic solvents were removed under vacuum and an aqueous layer was extracted using DCM. The final crude product was reprecipitated from DCM using pentane, yielding blue crystals (34 mg, 61 % yield).

^1^H NMR (401 MHz, CD2Cl2) δ 7.64 – 7.49 (m, 11H), 7.26 (d, *J* = 16.3 Hz, 2H), 7.01 – 6.92 (m, 4H), 6.66 (s, 2H), 4.03 (t, *J* = 5.2 Hz, 4H), 3.07 (t, *J* = 5.2 Hz, 4H), 1.64 (s, 4H), 1.46 (s, 6H). ^13^C NMR (101 MHz, CD2Cl2) δ 160.5, 152.9, 142.6, 136.2, 135.5, 133.6, 132.2, 130.1, 129.9, 129.5, 129.3, 129.0, 117.8, 117.3, 115.3, 114.9, 71.0, 41.9, 14.8. HR-MS (ESI+), m/z: [M+H]+, calcd for [C37H38O2N4BF2]^+^ 619.30571, found 619.30504. ^1^H NMR and ^13^C NMR of **BODIPY-NH2** are shown Figures S22 A, B respectively. HRMS spectrum of **BODIPY-NH2** is shown in Figure S23.

**BDP-NHAc:** To a solution of **BDP-NH2** (5 mg, 0.008 mmol, 1 eq.) in THF (2.0 ml), Et3N (15 mg, 0.14 mmol, 17 eq.) and CH3COCl (11 mg, 0.15 mmol, 18 eq) were added. The reaction mixture was stirred at r.t. for 2 h. Water was added and a product was extracted using DCM. The solvent was removed under reduced pressure and the crude product was 3 times reprecipitated from DCM using pentane, yielding blue crystals (4.5 mg, 79 %).

^1^H NMR (401 MHz, CD2Cl2) δ 7.78 – 7.46 (m, 10H), 7.40 – 7.31 (m, 3H), 7.26 (d, *J* = 16.3 Hz, 2H), 7.01 – 6.89 (m, 4H), 6.65 (s, 2H), 4.09 (t, *J* = 5.1 Hz, 4H), 3.65 (q, *J* = 5.4 Hz, 4H), 1.97 (s, 3H), 1.46 (s, 6H). ^13^C NMR (101 MHz, CDCl3) δ 160.0 (C), 152.9 (C), 142.7 (C), 139.1 (C), 136.0 (CH), 135.5 (C), 130.2 (C), 129.5 (CH), 129.3 (CH), 128.9 (CH), 117.8 (CH), 117.5 (CH), 115.2 (CH), 67.5 (CH2), 39.3 (CH2), 19.1 (CH3), 14.8 (CH3). HR-MS (ESI+), m/z: [M+H]+, calcd for [C41H41O4N4BF2Na]^+^ 725.30841, found 725.30811.

^1^H NMR and ^13^C NMR of **BODIPY-NHAc** are shown Figures S24 A, B respectively. HRMS spectrum of **BODIPY-NHAc** is shown in Figure S25.

## Supplementary Figure S1-S25

**Figure S1:**
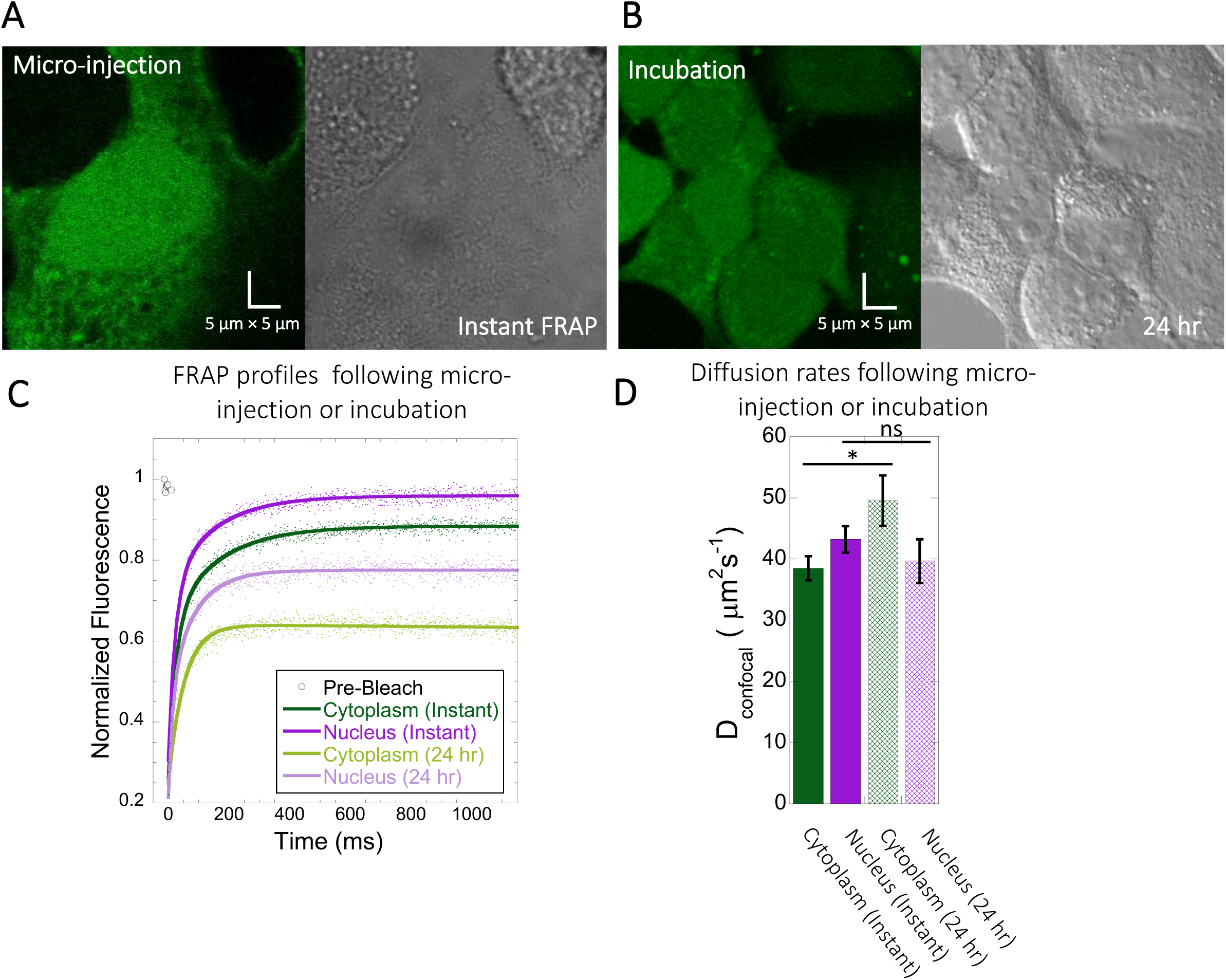
FRAP measurements for Fluorescein following different incubation times. (A) Micrographs of HeLa cells instantaneously after micro-injection *vs* (B) FRAP after 24 hours incubation. (C) Comparative merged FRAP profiles with fits (N= 30; R= 0.99 for each of the fits) and (D) their D_confocal_ values. Error bars represent SE calculated from fitting the FRAP progression curves averaged over at least 30 independent measurements.

**Figure S2:**
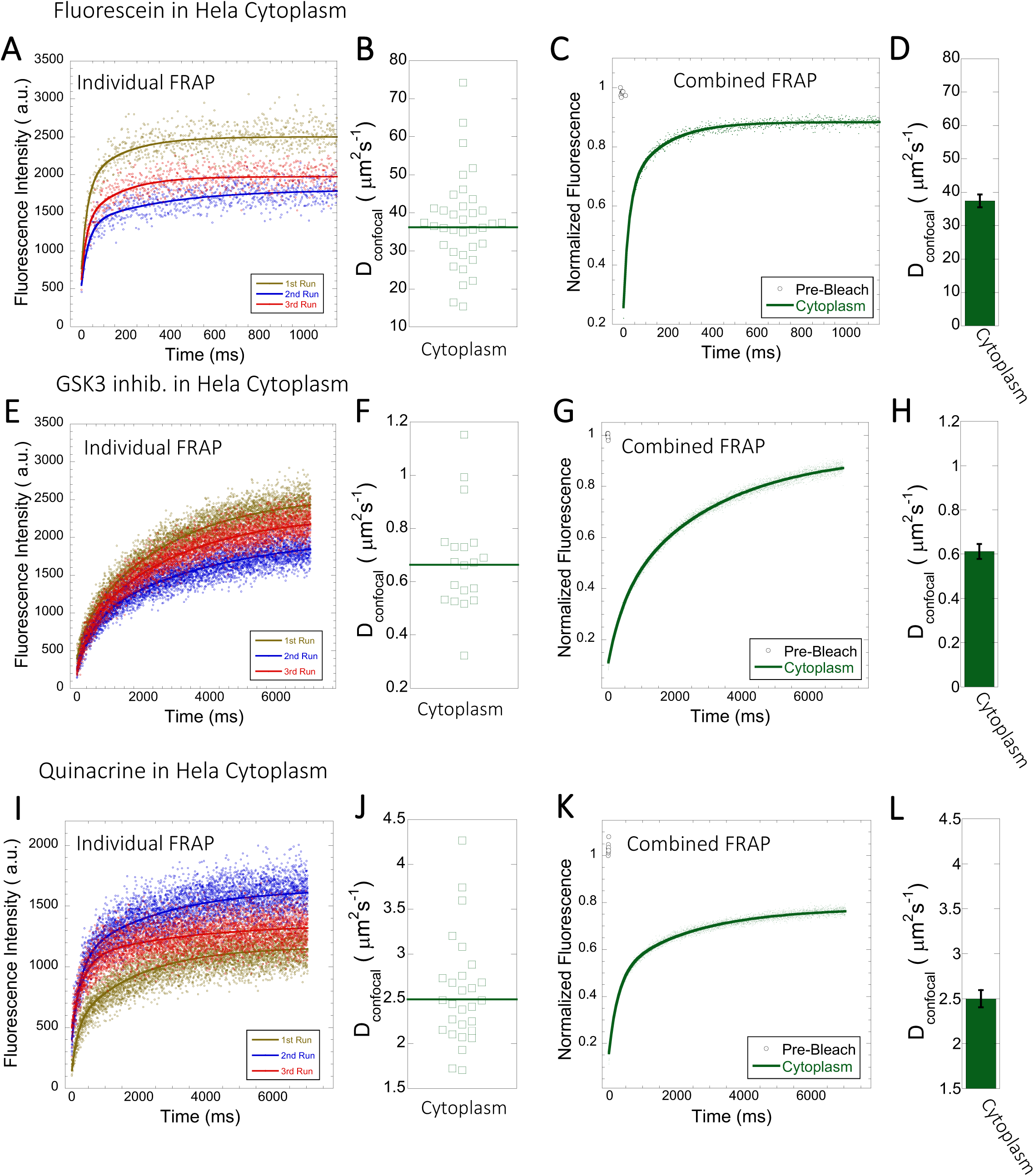
D_confocal_ values calculated from individual FRAP and merged FRAP curves. FRAP curves and D_confocal_ values for Fluorescein (A-D), GSK3 inhibitor (E-H) and Quinacrine (I-L) inside HeLa cell cytoplasm are shown. Three representative individual FRAP profiles with exponential fits with R= 0.80-0.90 for each of the compounds respectively are shown in (A), (E), and (I). Individual D_confocal_ values and their geometric means from individual FRAP curves are shown for Fluorescein((B), GSK3 inhibitor (F) and Quinacrine (J). (C), (G) and (K) show the merged FRAP curves over 30 independent measurements. This provides an improved fit (R=0.99). Panels (D), (H) and (L) show D_confocal_ values calculated from the merged FRAP profiles. Error bars represent SE calculated from fitting the FRAP progression curves.

**Figure S3:**
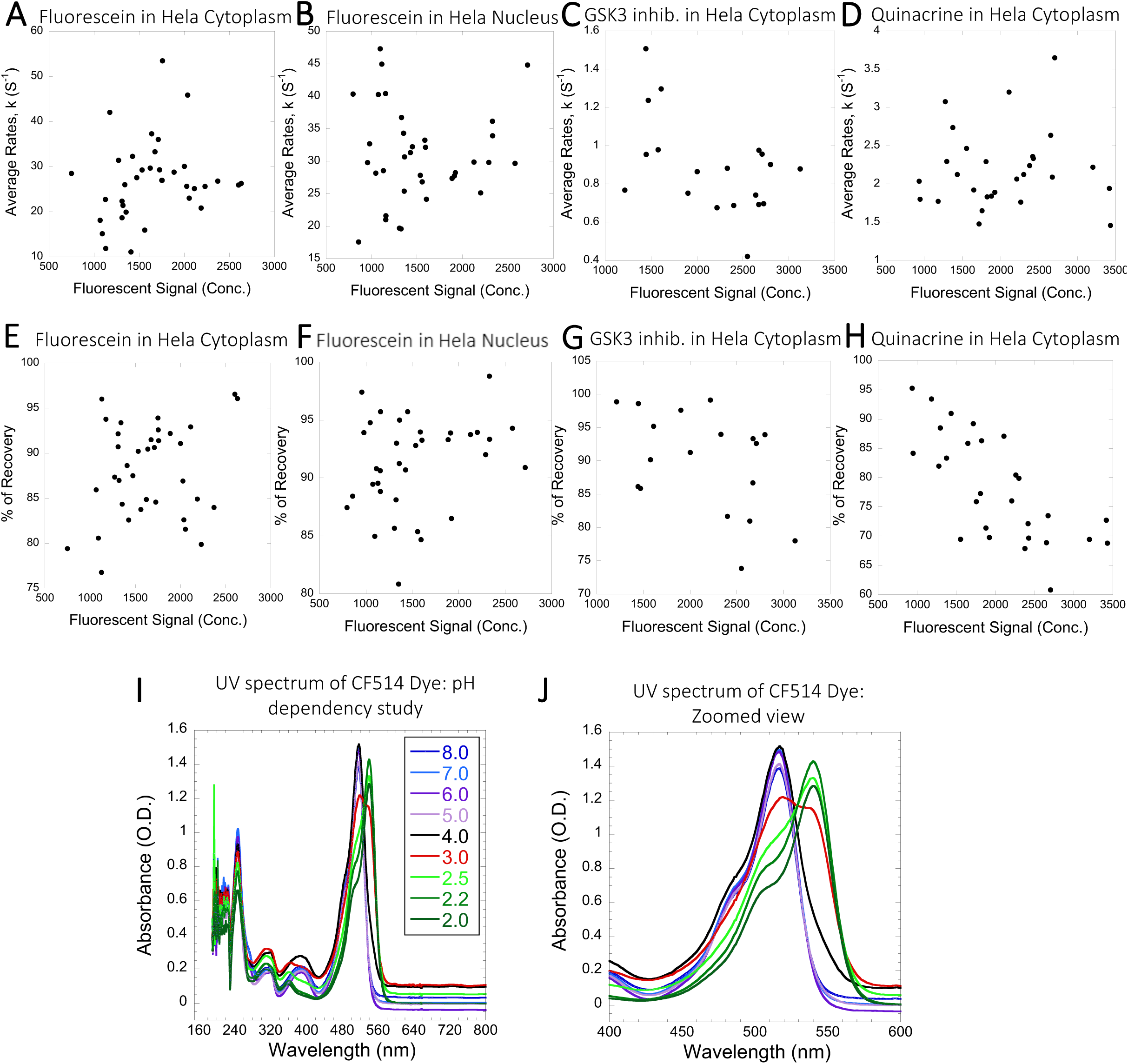
Diffusion rates and percent recovery as a function of drug concentration inside Hela Cells. The diffusion rates were calculated from individual FRAP progression curves. The drug concentration is linearly related to the fluorescence intensity. (A-D) show FRAP rates as a function of the fluorescence signal for Fluorescein in the cytoplasm. (B) Fluorescein in Nucleus, (C) GSK3 inhibitor in cytoplasm, (D) Quinacrine in cytoplasm. (E-H) are as (A-D) but the x-axis denotes percent recovery after FRAP. (I-J) Comparative UV spectrum of CF514 labelling dye in 20 mM sodium phosphate buffer at different pH values, from which a p*K*a value of 3 was calculated.

**Figure S4:**
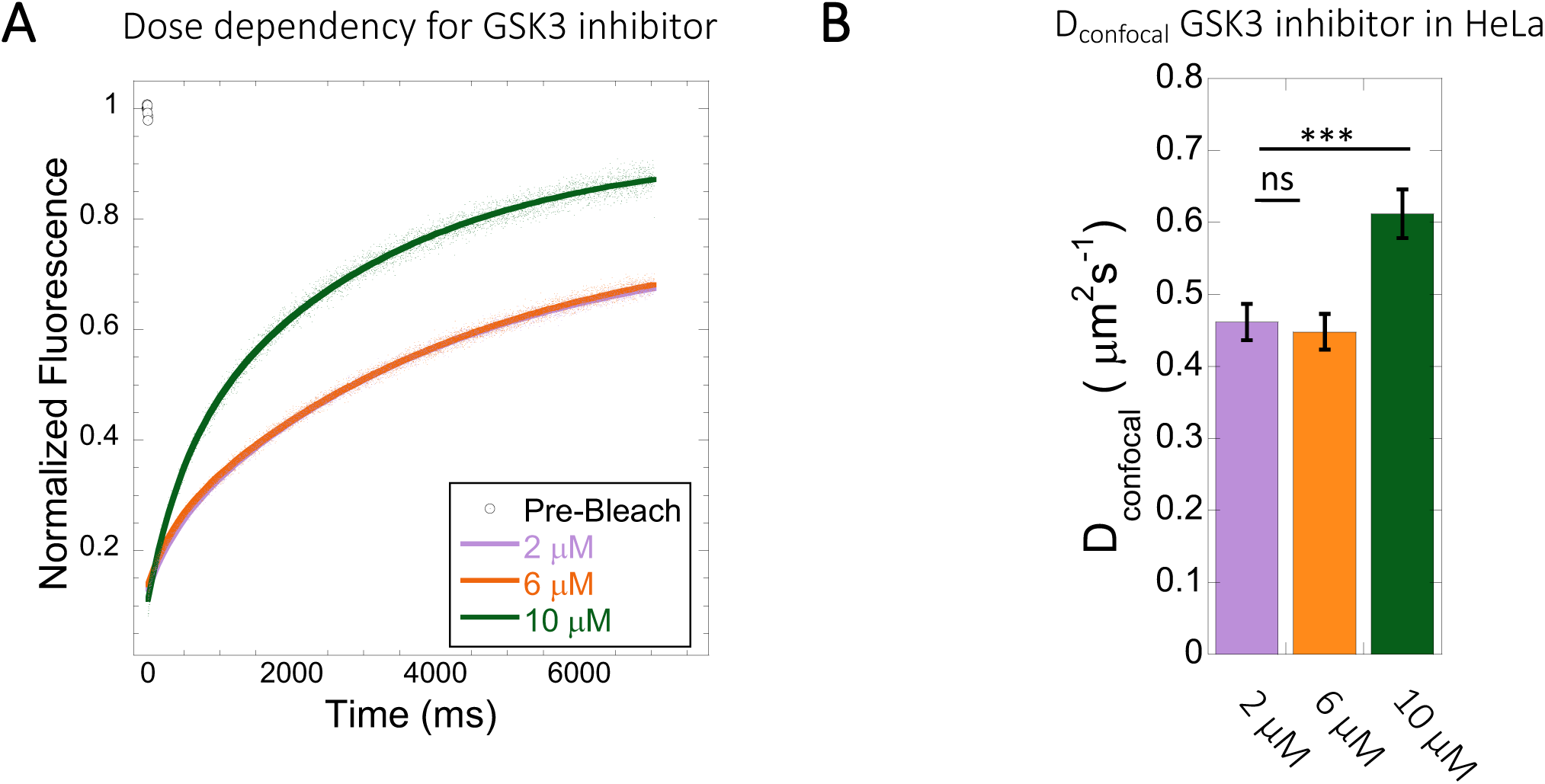
Diffusion rates at different doses of GSK3 inhibitor in HeLa cytoplasm. Effect of drug dosage (2-10 µM) on (A) averaged FRAP profiles with fits (N= 30; R= 0.99 for each of the fits) and (B) D_confocal_ values. Error bars represent SE calculated from fitting the FRAP progression curves, which are averaged over at least 30 independent measurements.

**Figure S5:**
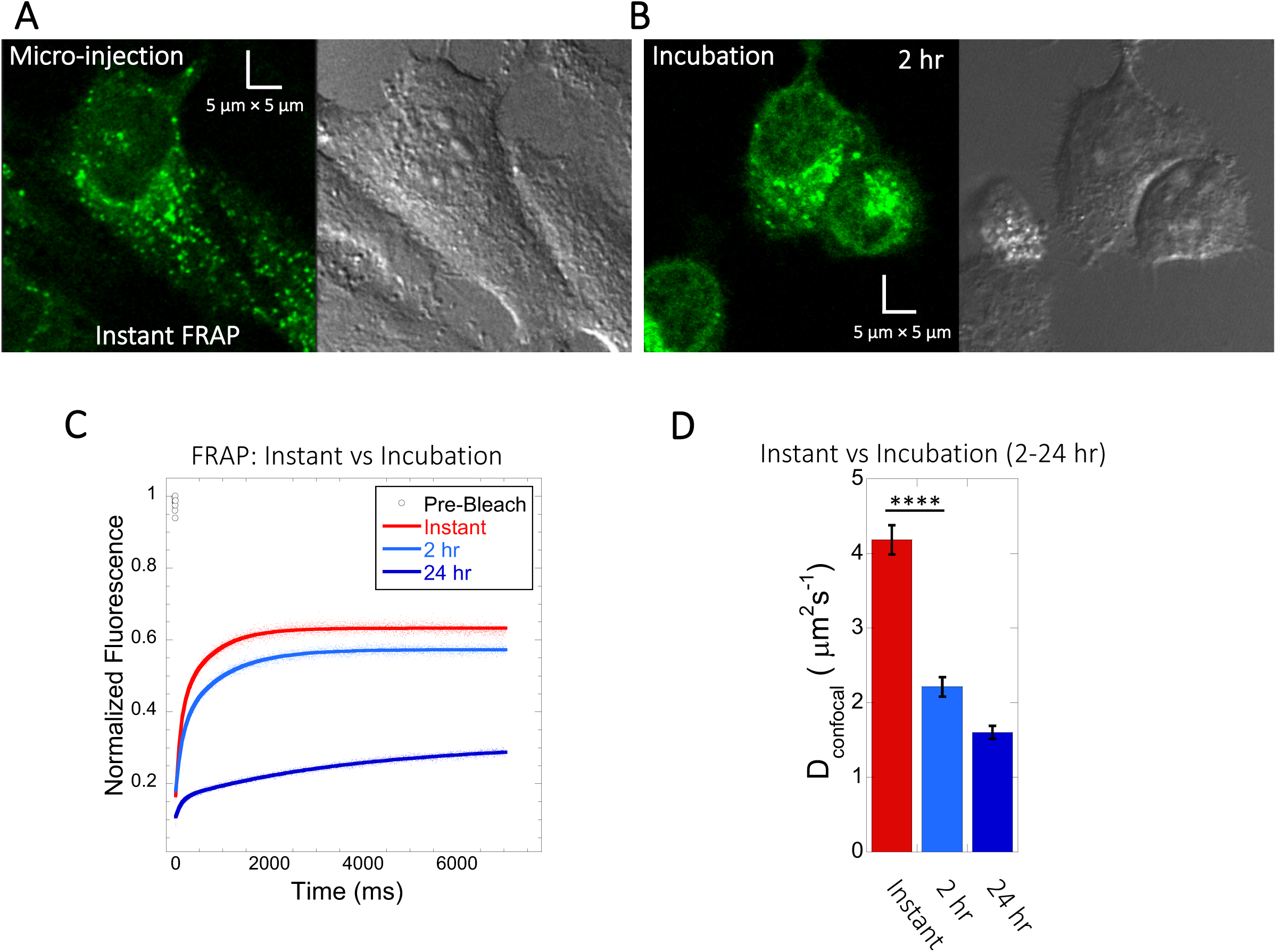
Quinacrine diffusion in HeLa after micro-injection or incubation. Comparison of FRAP immediately following micro-injection (instant FRAP) and FRAP following 2-24 hr incubation. HeLa cell micrographs following (A) micro-injection and (B) 2 hours of incubation. Comparative (C) averaged FRAP profiles with exponential curve fits (N= 30; R= 0.99 for each of the fits) and (D) calculated D_confocal_ values. Error bars represent SE calculated from fitting the FRAP progression curves, which are averaged over at least 30 independent measurements.

**Figure S6:**
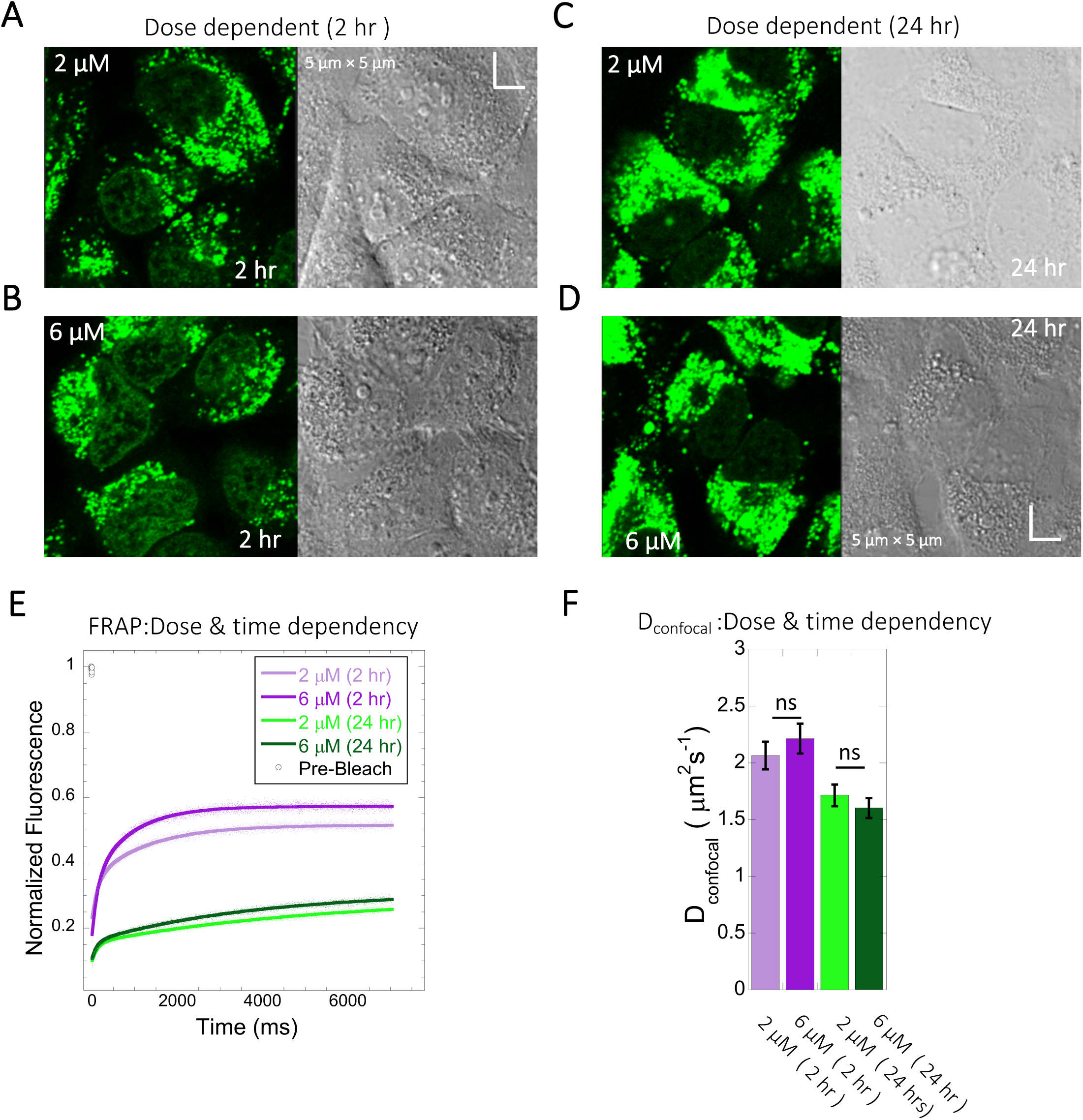
Dose and time dependency of Quinacrine diffusion in HeLa cells. HeLa cell micrographs after 2 hours of incubation with (A) 2 µM and (B) 6 µM of Quinacrine. (C) and (D) are micrographs of HeLa cells after 24 hours of incubation with 2 µM and 6 µM Quinacrine respectively. (E) Effect of drug dose and time dependency on comparative FRAP profiles, with fits (N= 30; R= 0.99 for each of the fits) and (F) Calculated D_confocal_ values from the fits in (E). Error bars represent SE calculated from fitting the FRAP progression curves, which are averaged over at least 30 independent measurements.

**Figure S7:**
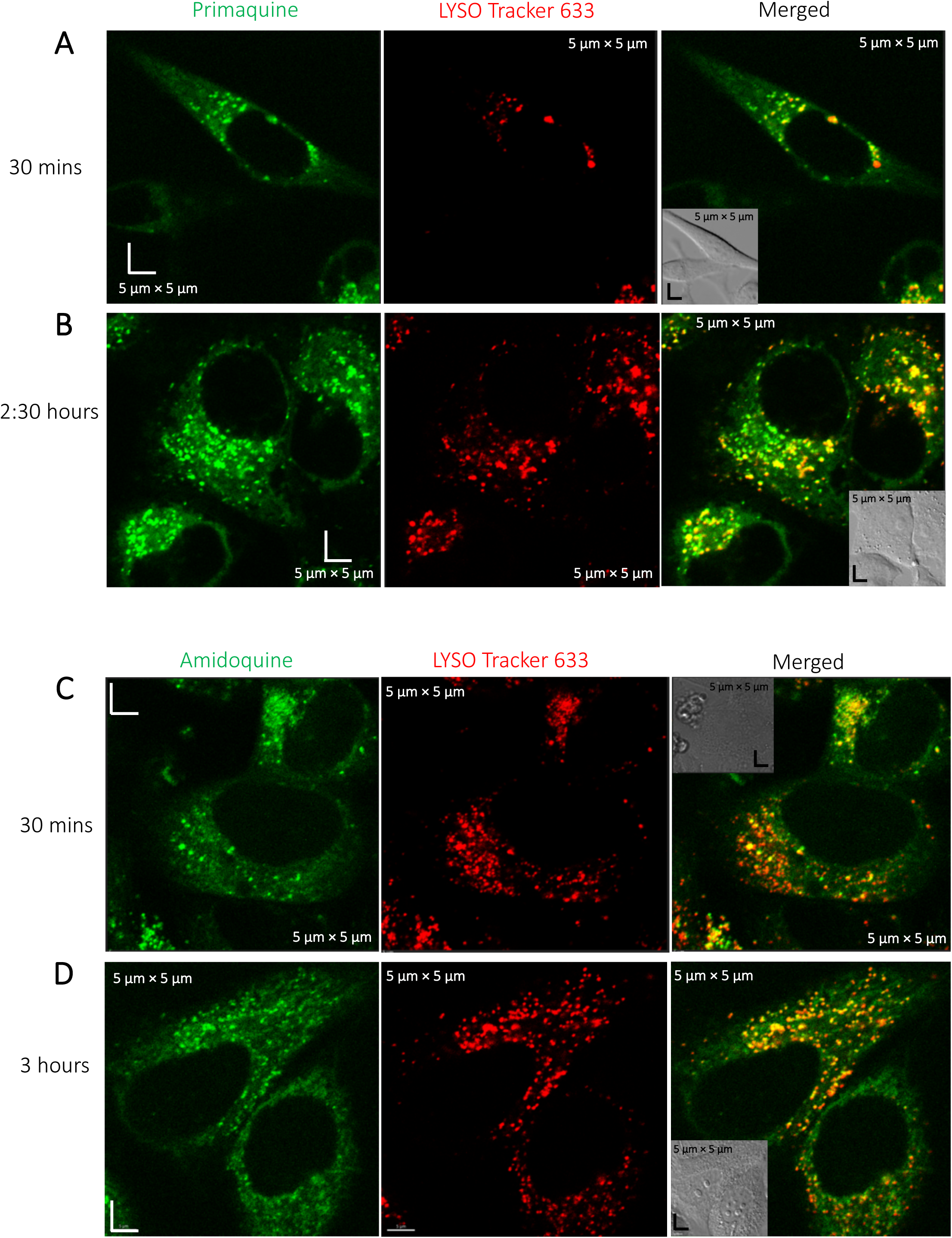
Colocalization of Primaquine and Amidoqiuine with Lyso-tracker. Colocalization of Primaquine (A-B) and Amidoquine (C-D) with Lyso Tracker 633 in HeLa cells. Time dependent measurements show higher aggregation of both drugs in the lysosomes over time.

**Figure S8:**
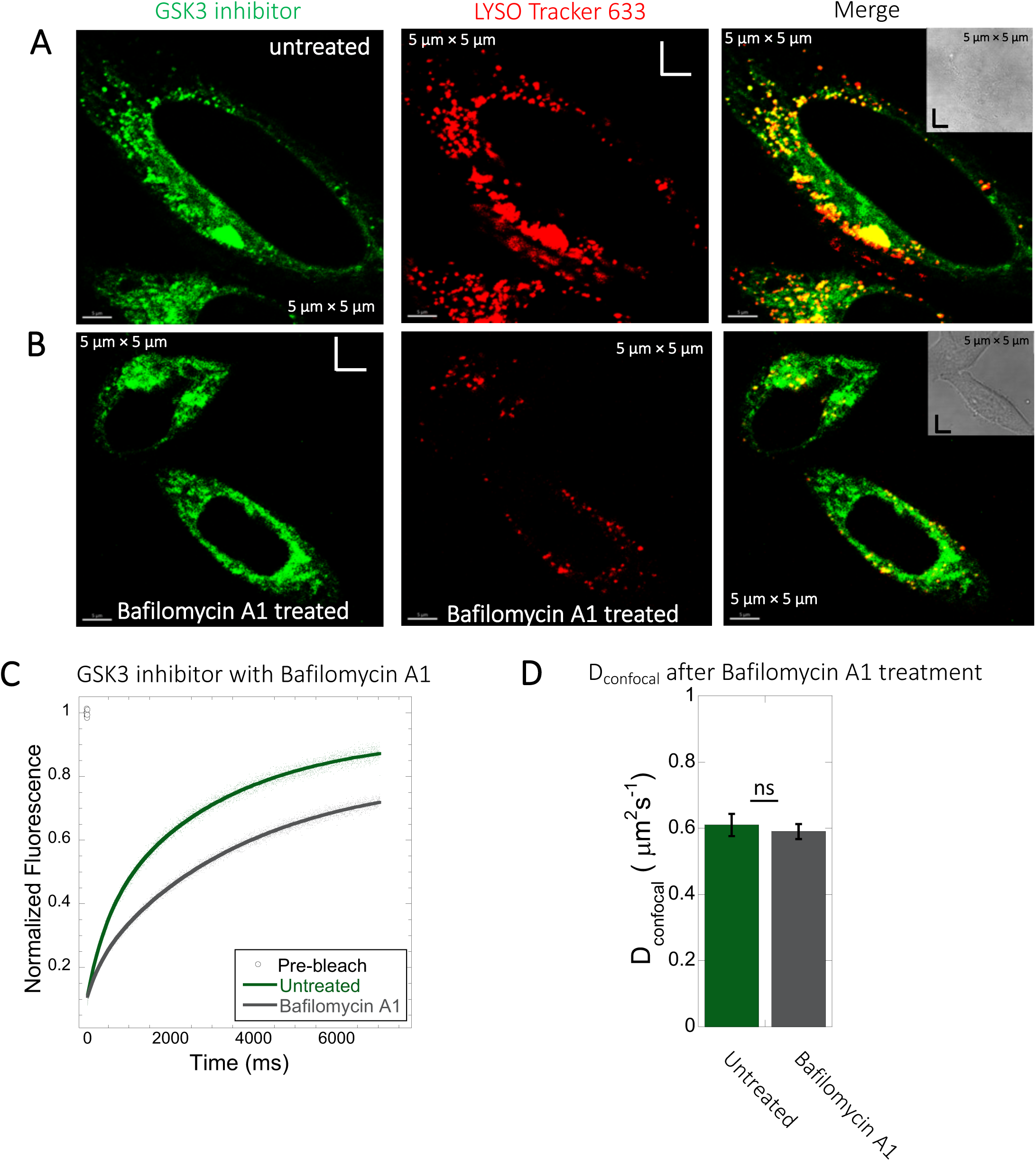
Inhibition of lysosomal accumulation of GSK3 inhibitor by Bafilomycin A1. (A-B) Colocalization of GSK3 inhibitor with Lyso Tracker 633 in HeLa cells. (A) Untreated and (B) treated with 100 nM Bafilomycin A1 and corresponding colocalizations are shown. (C) Comparative averaged FRAP profiles with fits (N= 30; R= 0.99 for each of the fits) and (D) D_confocal_ diffusion values with and without the Bafilomycin A1 treatment. Error bars represent SE calculated from fitting the FRAP progression curves, which are averaged over at least 30 independent measurements.

**Figure S9:**
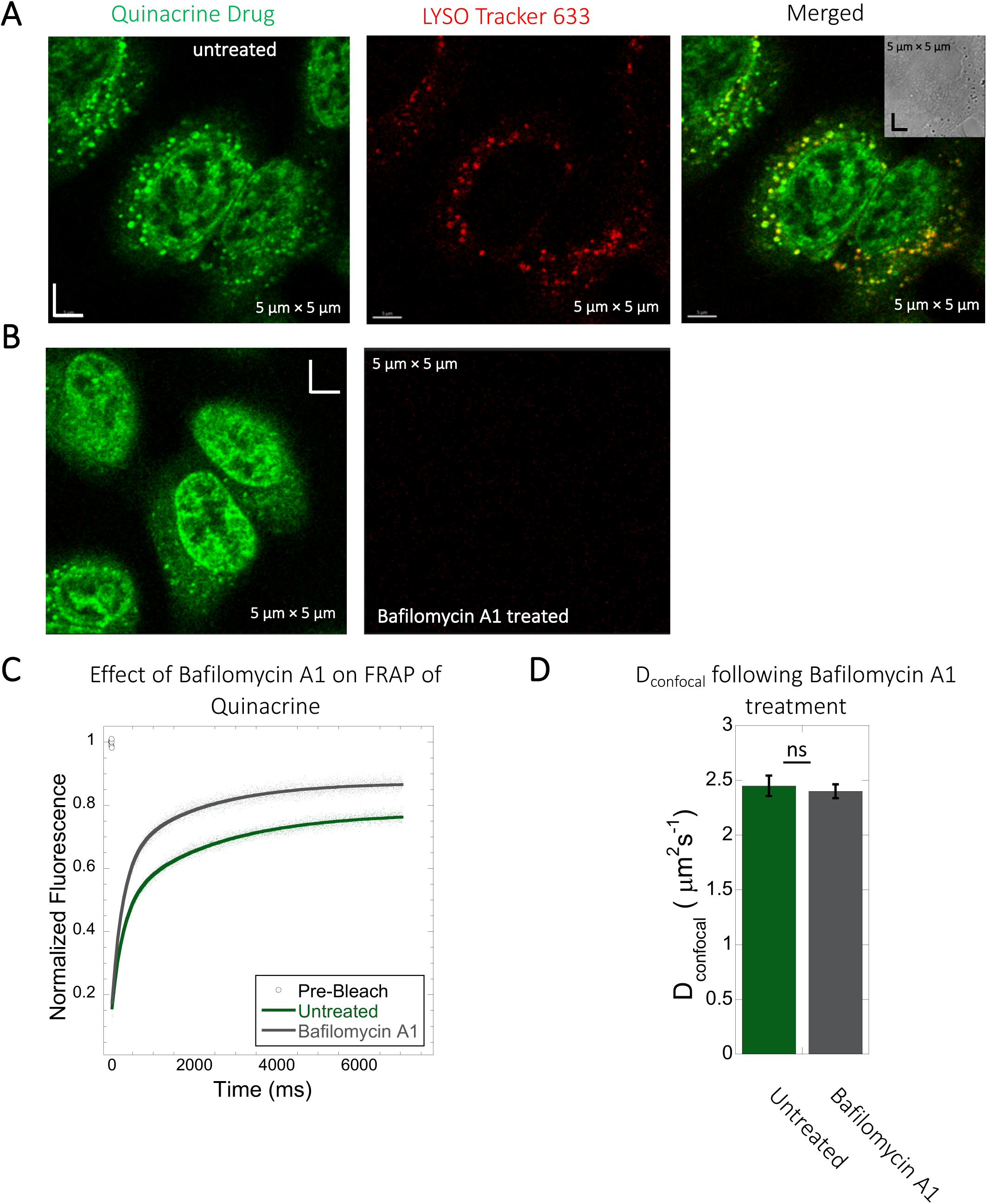
Inhibition of lysosomal accumulation of Quinacrine by Bafilomycin A1. (A-B) Colocalization of Quinacrine with Lyso Tracker 633 in HeLa cells. HeLa cells (A) untreated and (B) treated with 100 nM Bafilomycin A1 and corresponding colocalizations. (C) Comparative averaged FRAP profiles with fits (N= 30; R= 0.99 for each of the fits) and (D) D_confocal_ values with and without the Bafilomycin A1 treatment. Error bars represent SE calculated from fitting the FRAP progression curves, which are averaged over at least 30 independent measurements.

**Figure S10:**
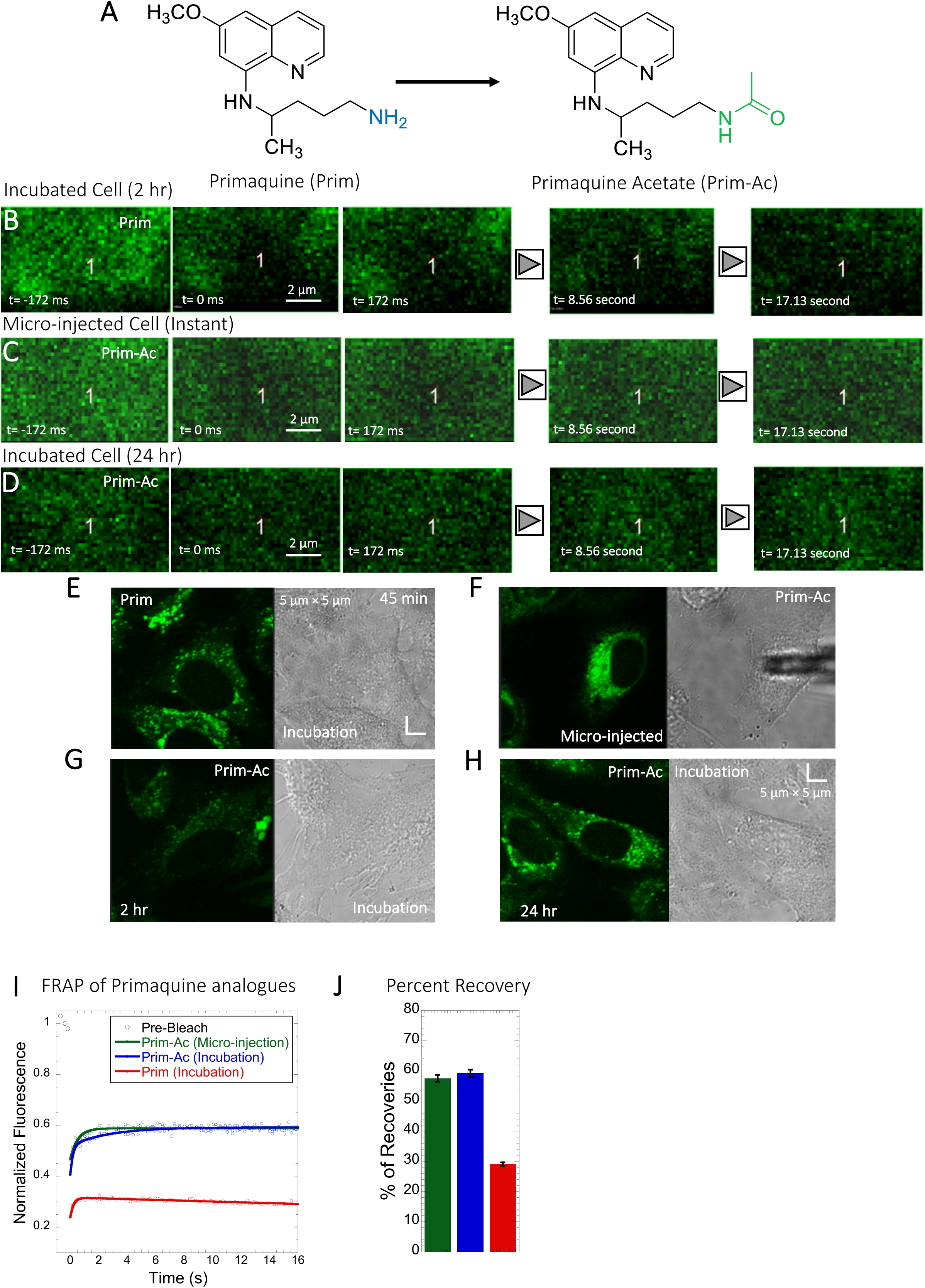
Comparative FRAP of Primaquine analogues in HeLa cell cytoplasm. (A) Structures of Primaquine analogues with p*K*a values. Comparative (B-D) XY FRAP micrographs with time lapses in HeLa cell. (E-H) Micrographs of treated HeLa cells either with micro-injection or after drug incubations with Prim or Prim-Ac. Comparative (I) XY-FRAP recovery profiles with fits (N= 20; R= 0.98 for each of the fits) and (J) percentage of recoveries for Primaquine analogues in HeLa cytoplasm (N= 20).

**Figure S11:**
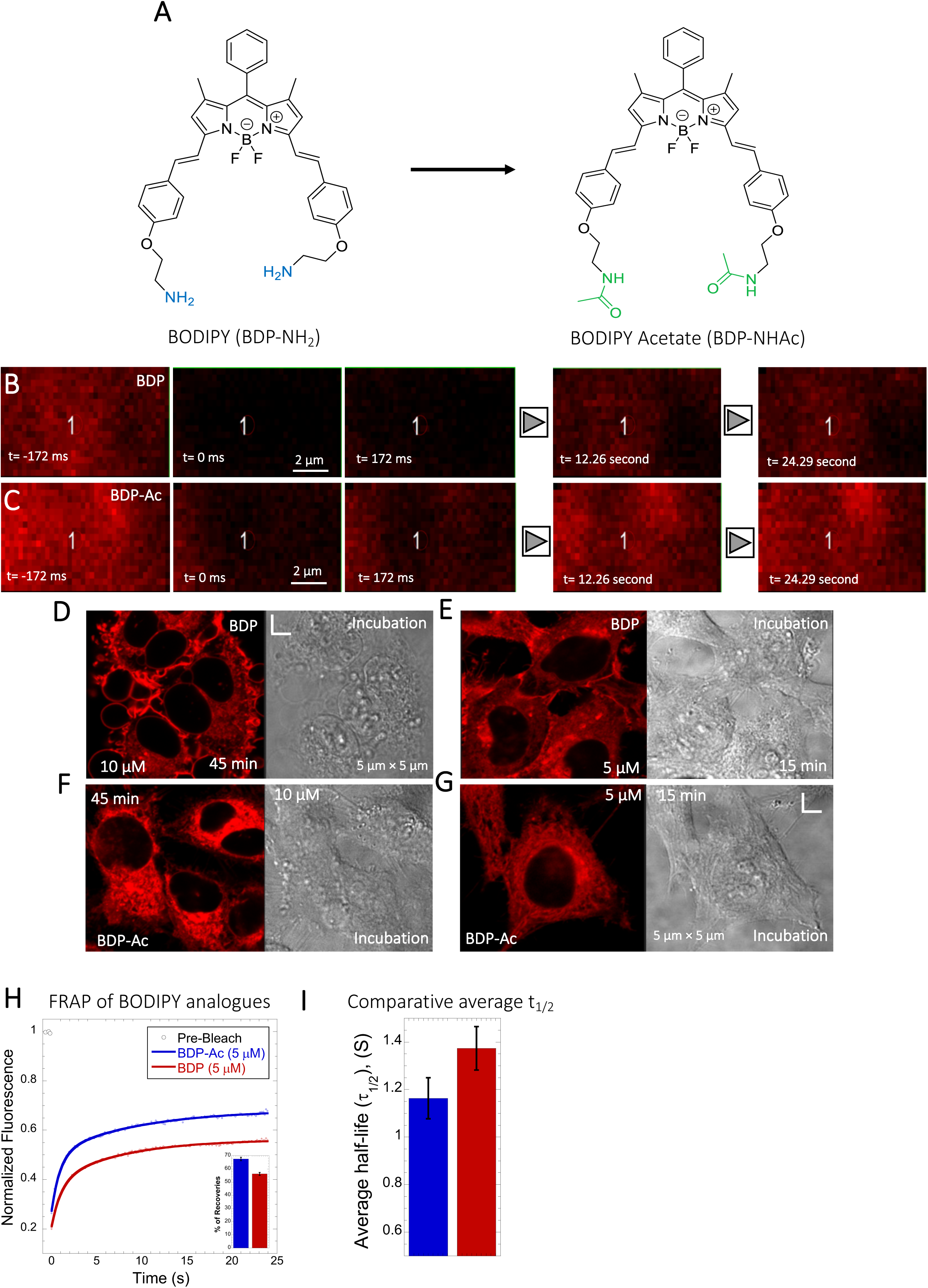
Comparative FRAP of BODIPY analogues in HeLa cell cytoplasm. (A) Structures of BODIPY analogues. (B-C) Comparative XY FRAP micrographs with time lapses in HeLa cell cytoplasm. (D-G) Micrographs of treated HeLa cells after incubations (with BDP/BDP-Ac) and, Comparative (H) averaged XY-FRAP recovery profiles with fits (N= 30; R= 0.99 for each of the fits), (I) averaged half-life and percentage of recoveries for BODIPY analogues in HeLa cytoplasm (H) (N= 30).

**Figure S12:**
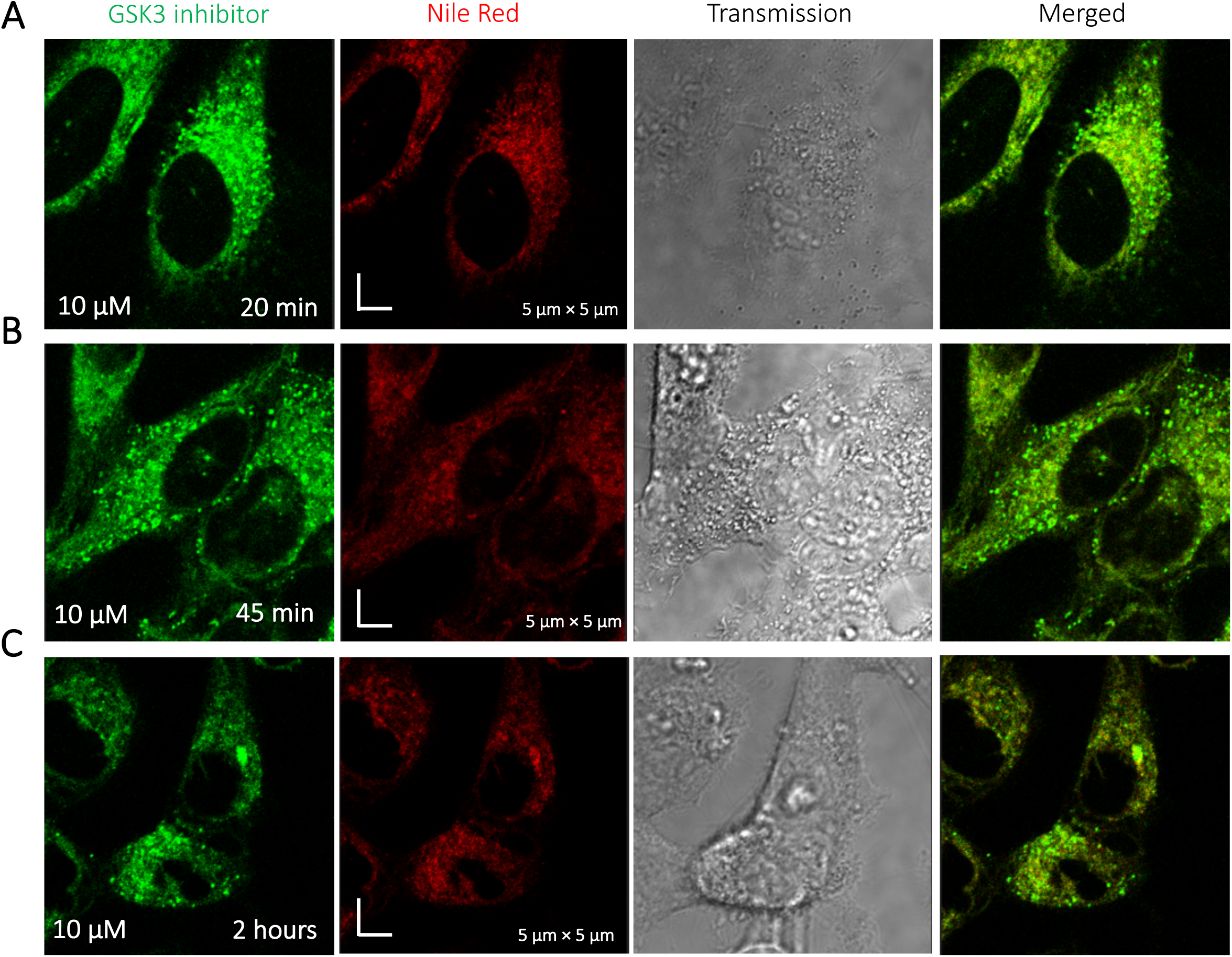
Time dependent colocalization of GSK3 inhibitor and Nile Red in HeLa cell cytoplasm. Panels (A-C) show Nile red accumulating in lipid droplets, while GSK3 inhibitor does not colocalize.

**Figure S13:**
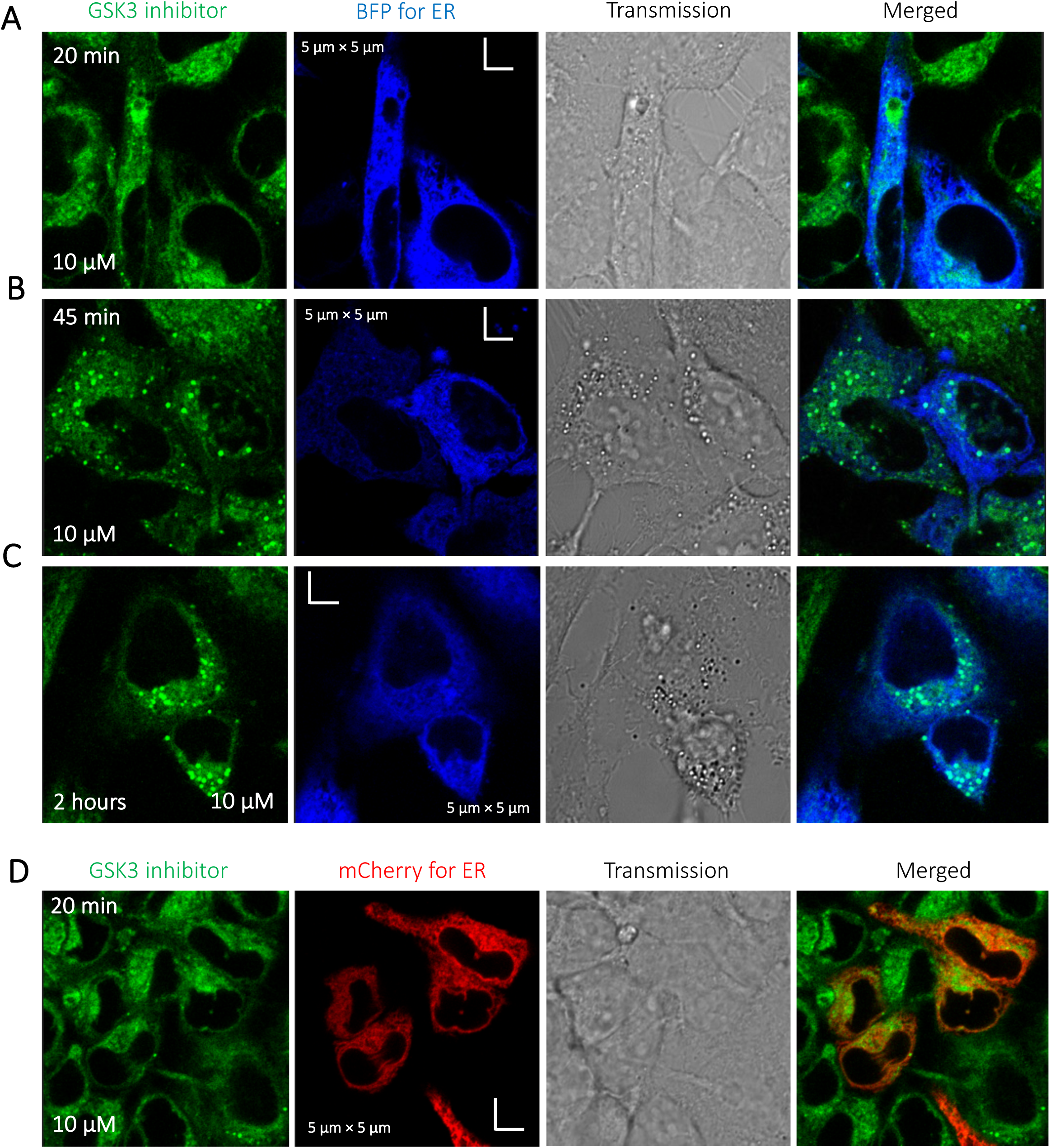
Time dependent colocalization of GSK3 inhibitor and ER marker in HeLa cell cytoplasm. Panels (A-D) show BFP/mCherry fused antibody marker accumulating in ER, but GSK3 inhibitor does not colocalize. HeLa cells were first transiently transfected with BFP/mCherry fused antibody markers, then treated with GSK3 inhibitor.

**Figure S14:**
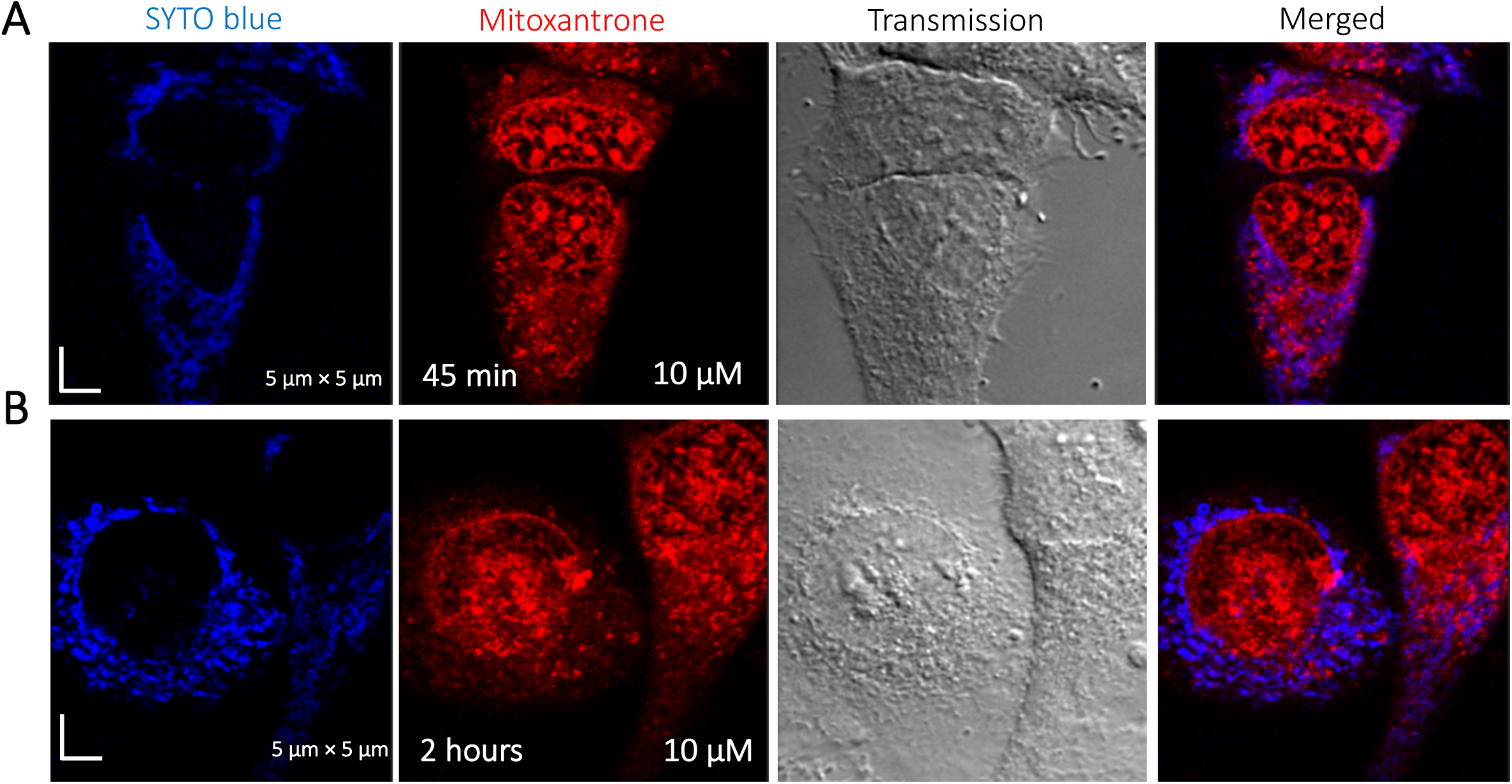
Time dependent colocalization study of Mitoxantrone and SYTO blue marker in HeLa cells. Panels (A-B) show SYTO blue marker accumulating in nucleic acids, but Mitoxantrone is not colocalized. HeLa cells were first treated with SYTO blue, then treated with Mitoxantrone.

**Figure S15:**
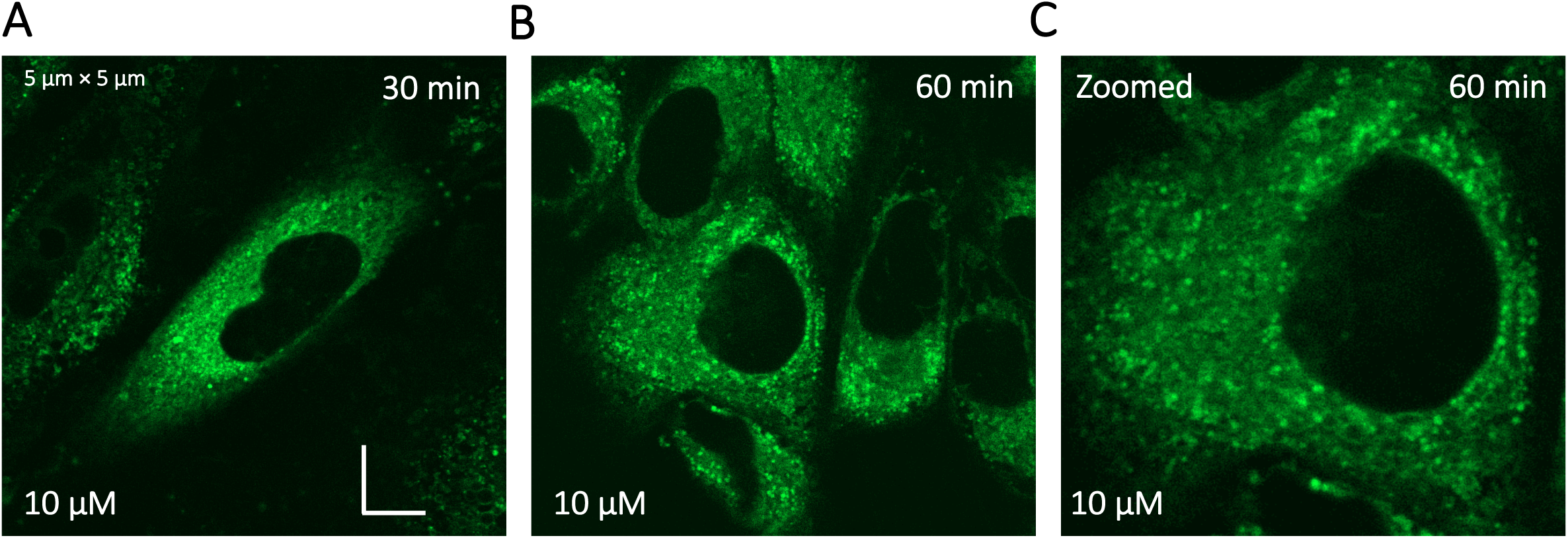
Super resolution images of GSK3 inhibitor treated HeLa cells. Micrographs were taken using a Zeiss LSM900 microscope.

**Figure S16:**
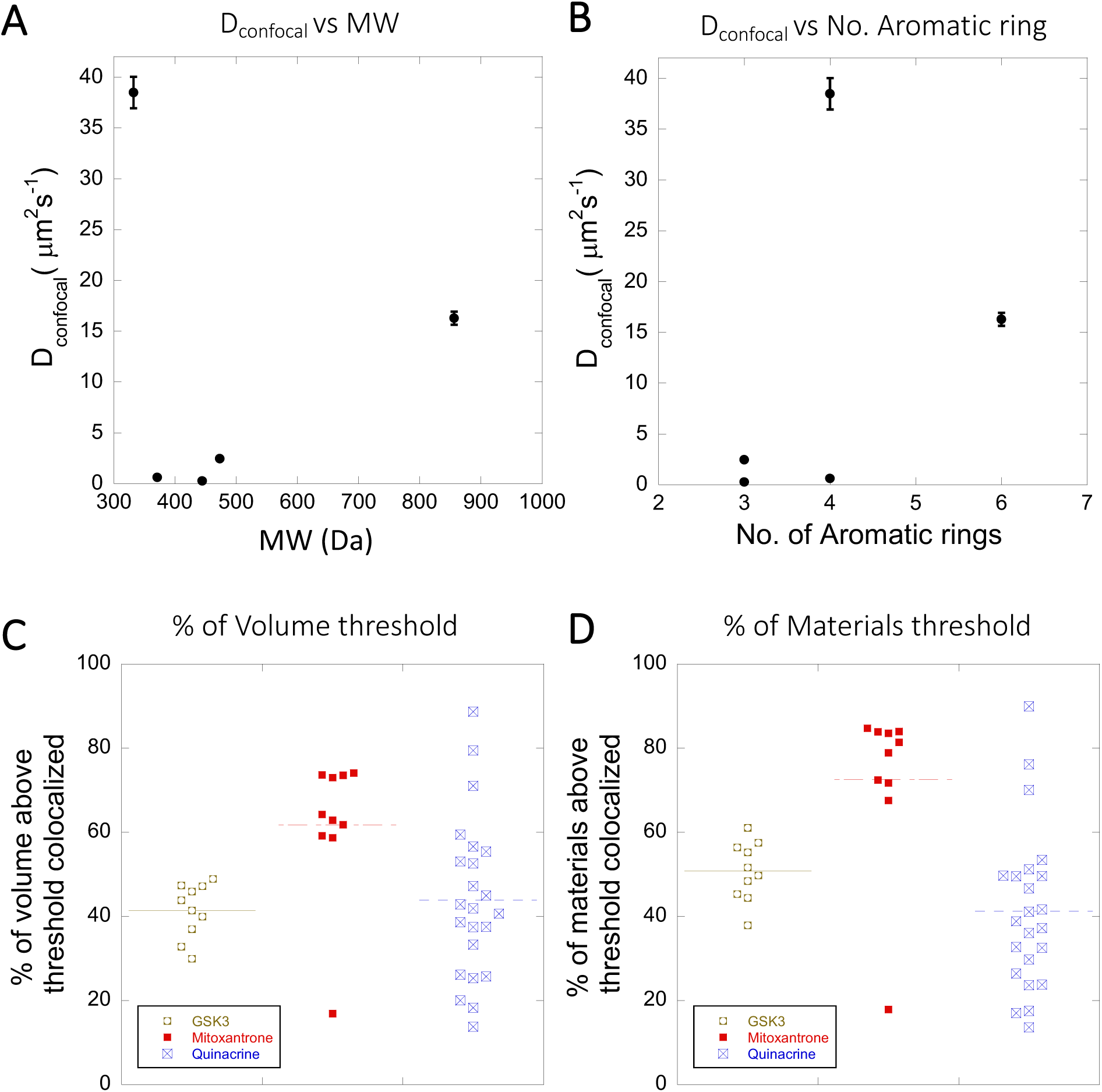
Dependency of D_confocal_ rates on MW, number of aromatic rings and colocalization within Lysosomes. (A) Dependency of D_confocal_ on molecular weight (MW). (B) Dependency of D_confocal_ on the number of aromatic rings in the drugs. (C-D) Comparison of percentage of colocalization among GSK3 inhibitor vs Mitoxantrone vs Quinacrine in acidic lysosomes are shown. Each dot represents 4-5 stained cells in a single frame acquisition. Colocalization study for GSK3 inhibitor and Quinacrine was done with commercial Lyso tracker-633, and for Mitoxantrone using Quinacrine itself used as a Lyso tracker dye.

**Figure S17:**
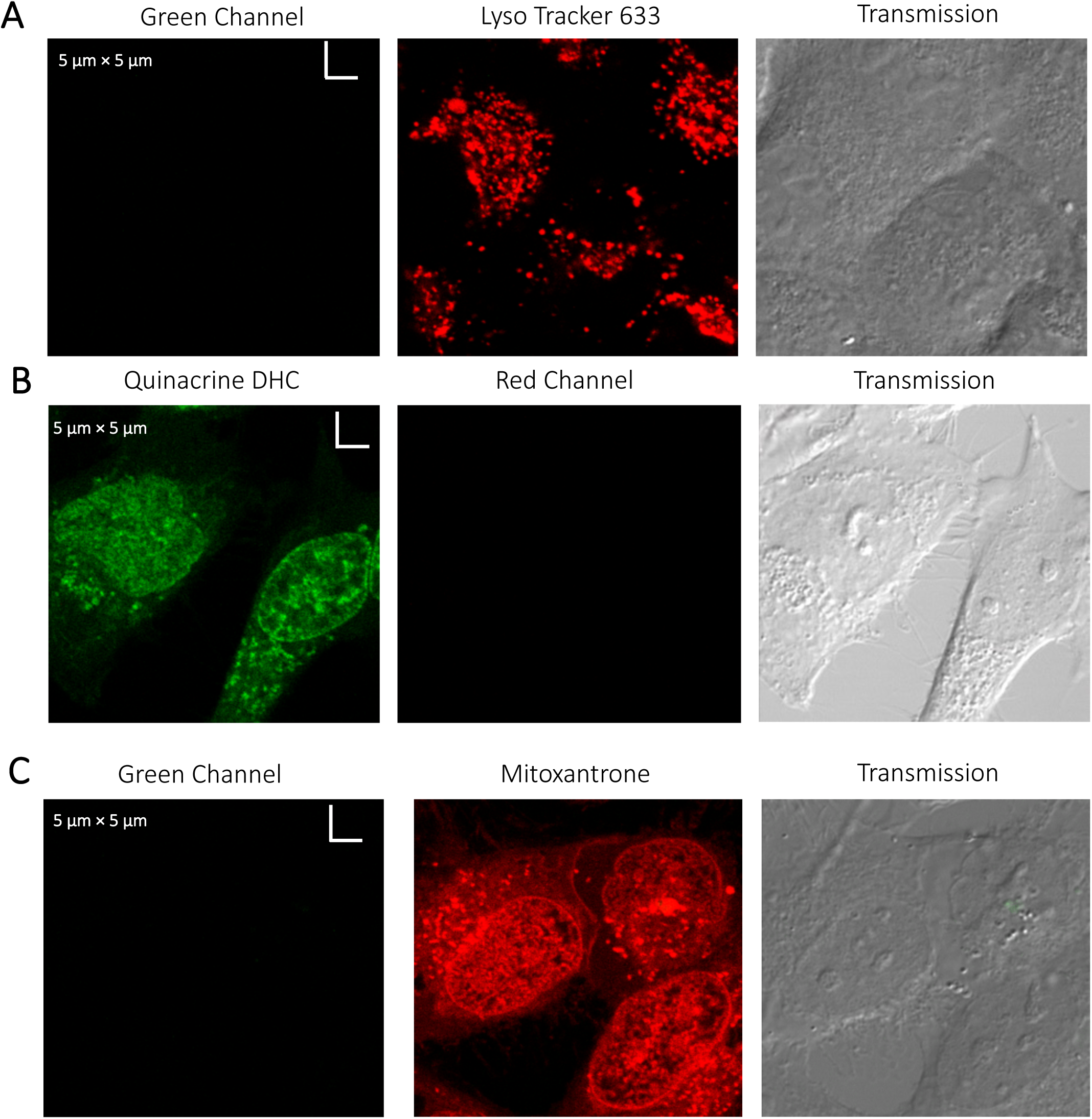
Evaluating channel leakage. Control experiments showing that channel leakage did not occur during colocalization studies. (A) Lyso-Tracker only, (B) Quinacrine only and (C) Mitoxantrone only.

**Figure S18:**
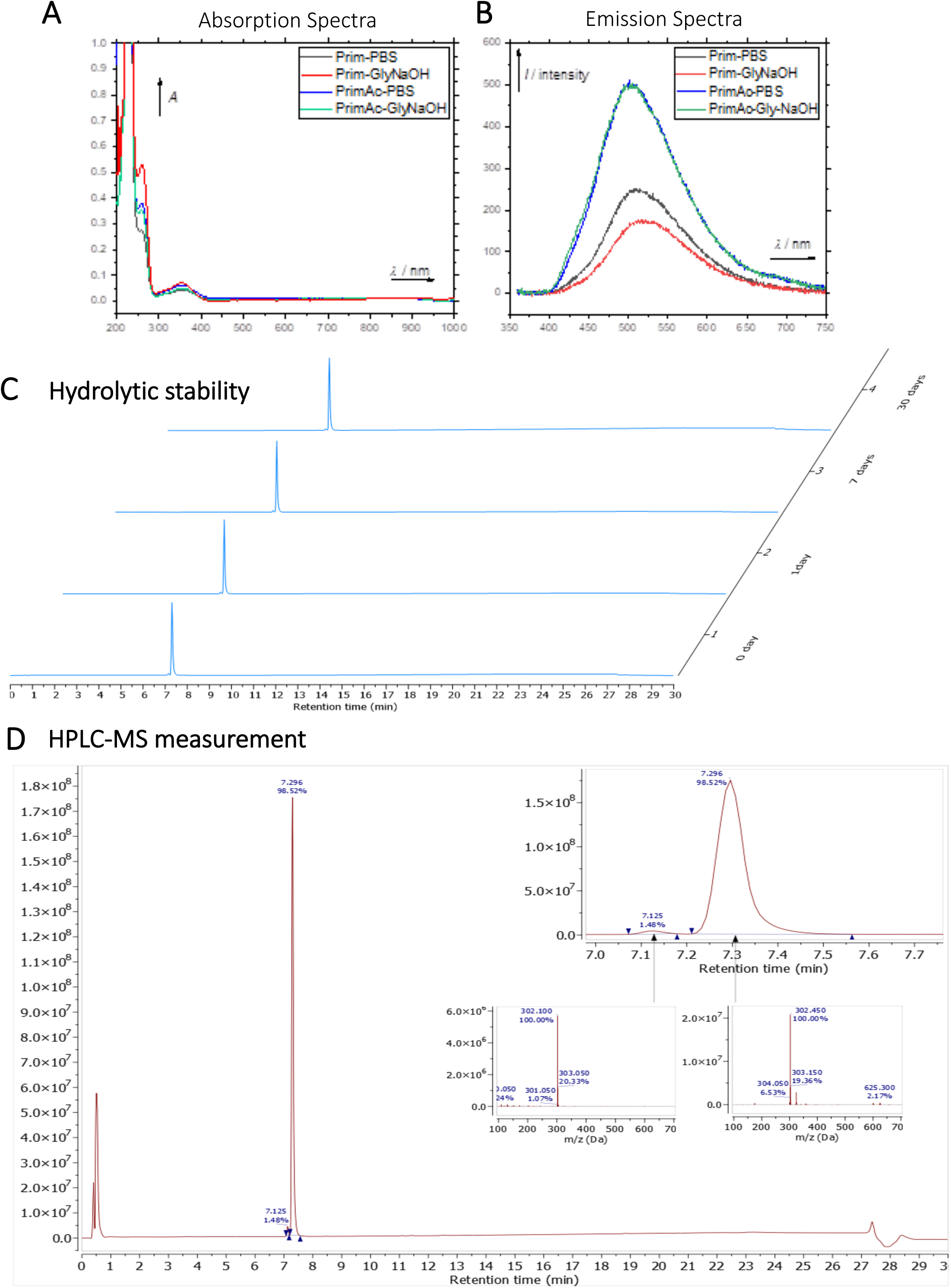
Characterization of Primaquine (Prim) and N-acetylated Primaquine (PrimAc). (A) Absorption (c ≈ 1 · 10-3) and (B) emission (c ≈ 3 · 10-4) spectra. Both experiments were measured in the mixture of 10% DMSO in PBS or in Glycine/NaOH buffer. (C) Measurement of hydrolytic stability by HPLC. (D) HPLC-MS measurement of PrimAc are shown. According to HPLC-MS data the peak at 7.125 corresponds to 302 m/z (ESI +) so as the main peak at 7.296. The molar mass of PrimAc corresponds to the measured value.

**Figure S19:**
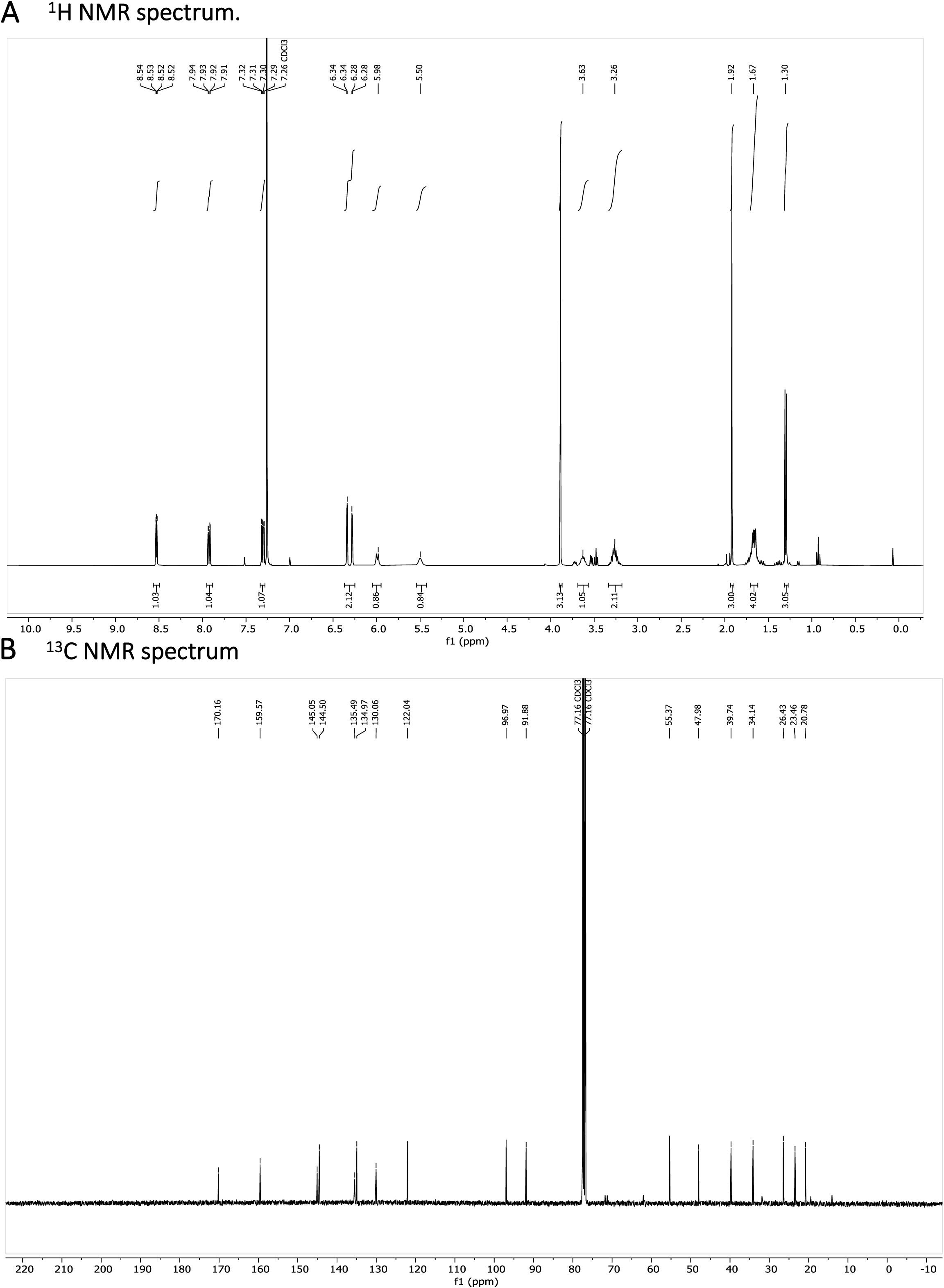
N-acetylated Primaquine. (A) ^1^H NMR spectrum is shown. The presence of impurities is caused by the degradation in the used solvent. (B) ^13^C NMR spectrum is shown.

**Figure S20:**
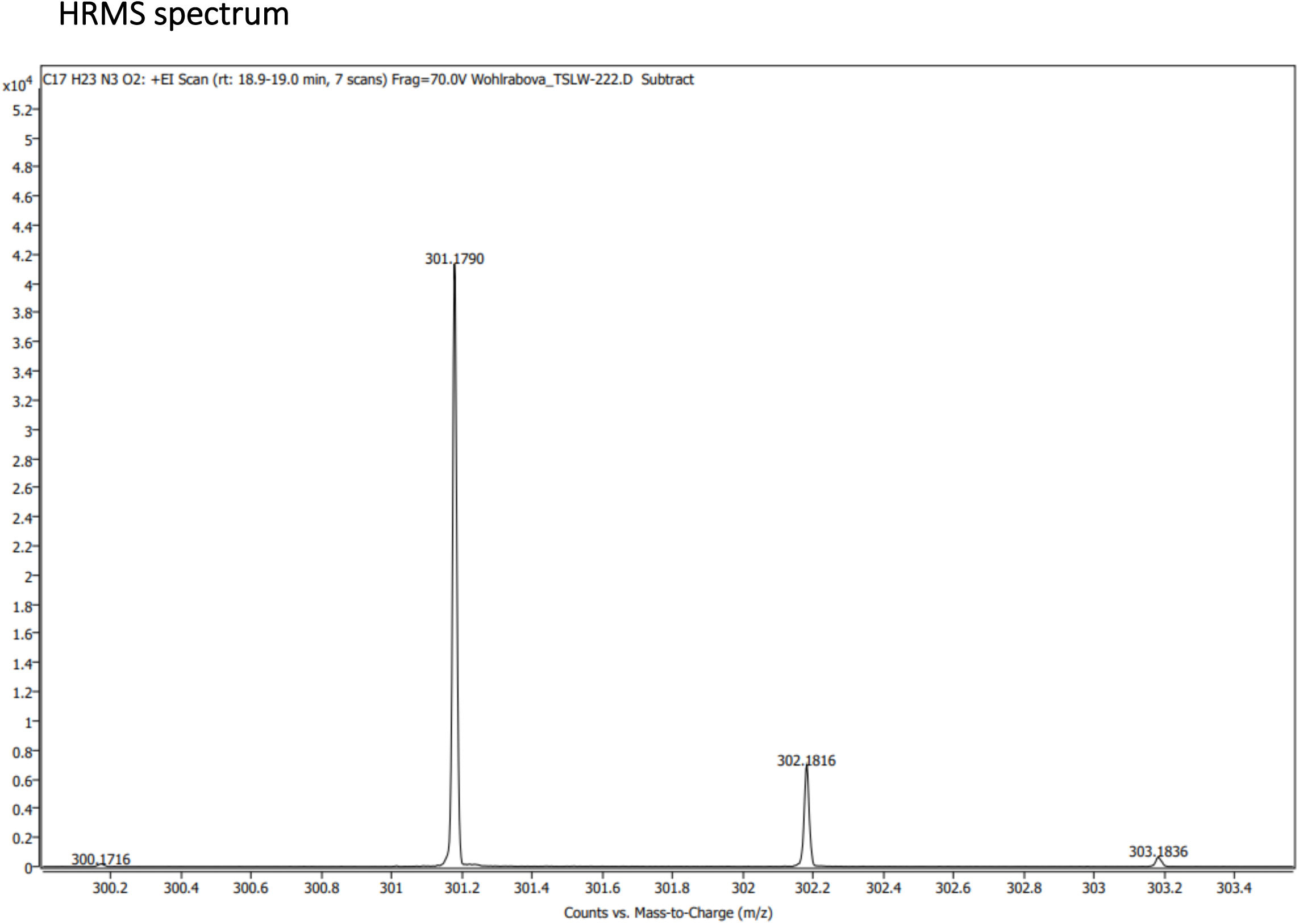
N-acetylated Primaquine. HRMS spectrum (+EI) is shown

**Figure S21:**
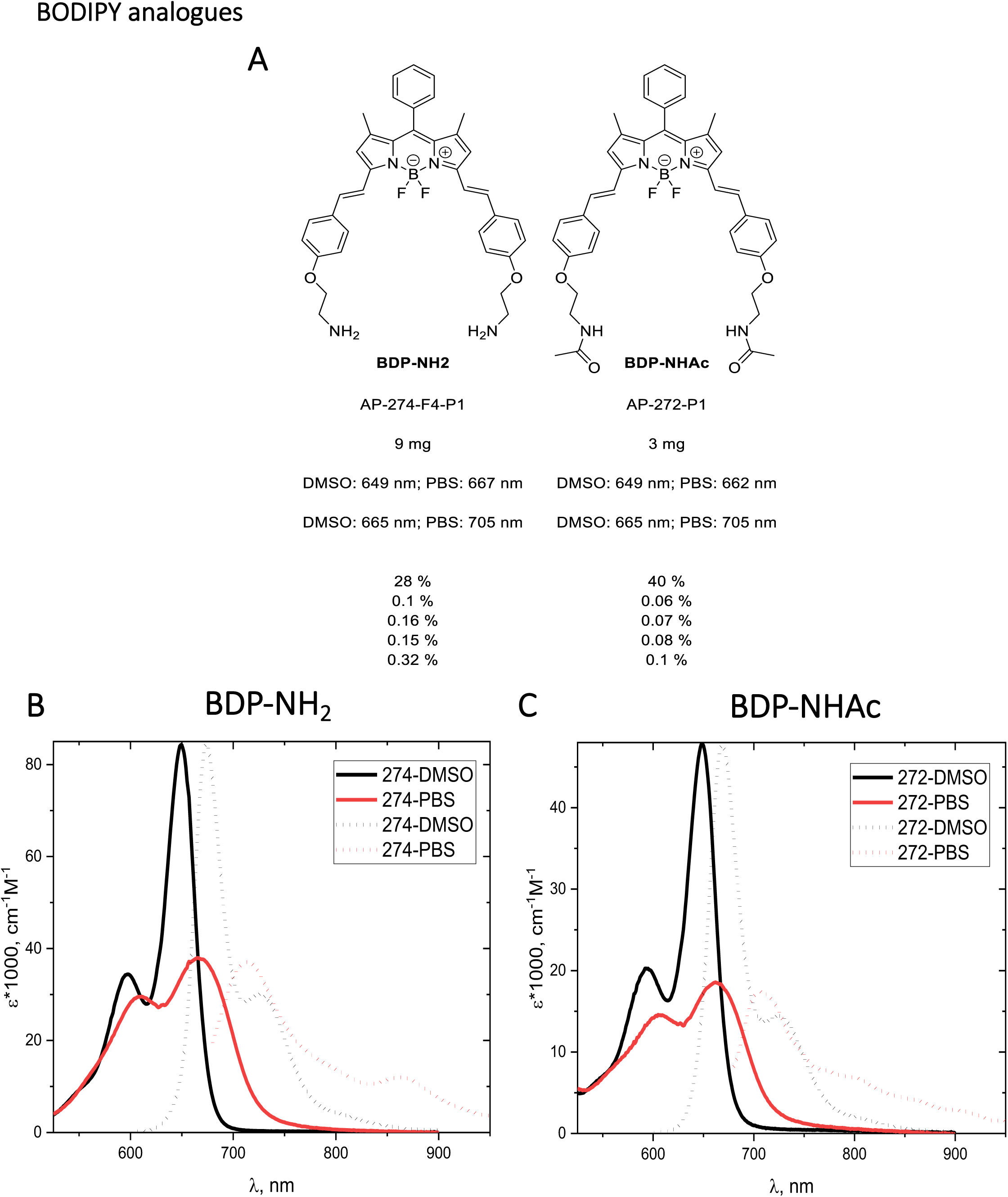
(A) Structure and photo-physical properties of BODIPY analogues. (B) Absorption (full line) and emission spectra (dashed line) of BDP-NH_2_ in DMSO (black) and PBS (red) are shown. (C) Absorption (full line) and emission spectra (dashed line) of BDP-NHAc in DMSO (black) and PBS (red) are shown.

**Figure S22:**
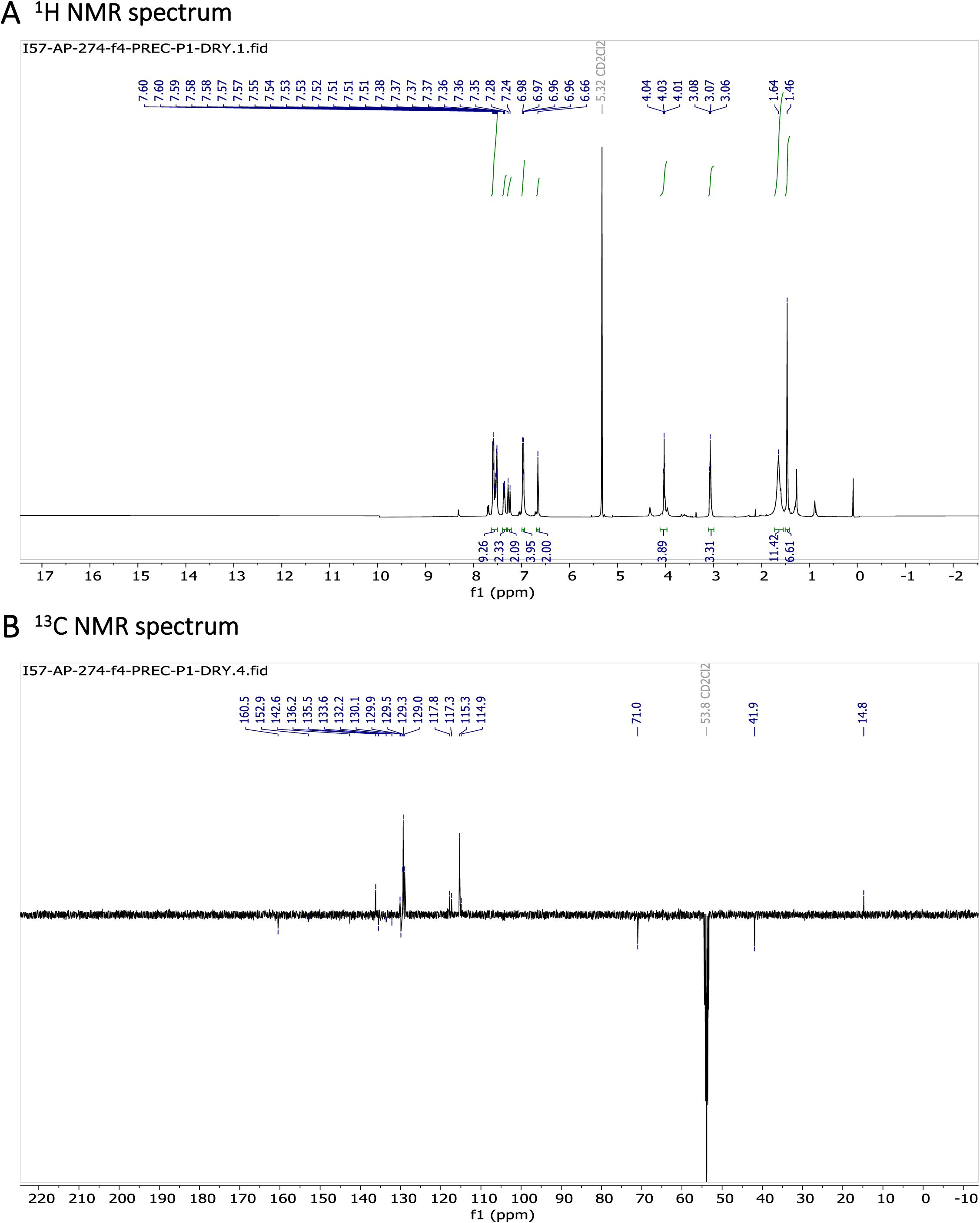
Characterization of BODIPY-NH_2_. (A) ^1^H NMR spectrum and (B) ^13^C NMR spectrum are shown.

**Figure S23:**
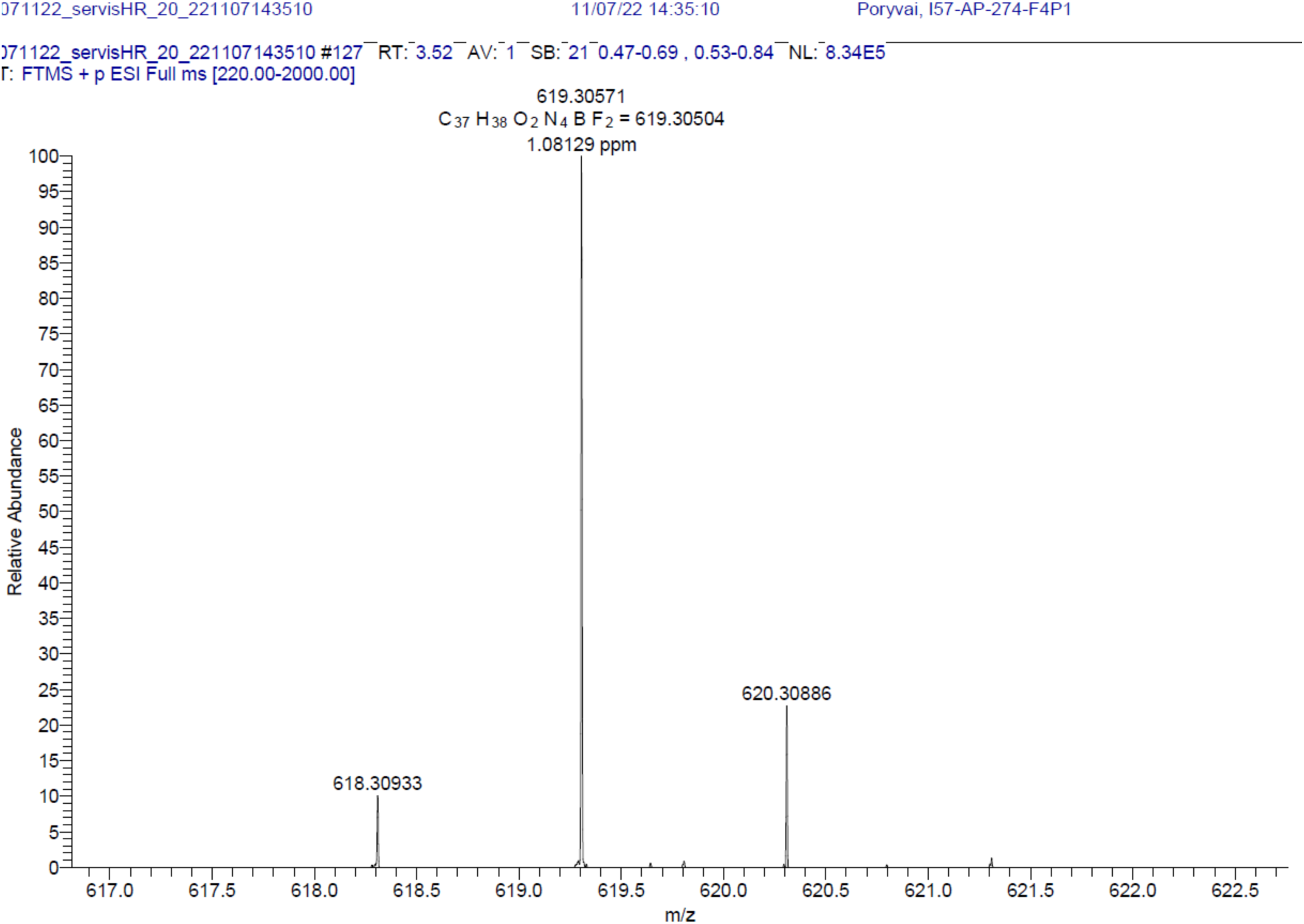
BODIPY-NH_2_. HRMS spectrum (+EI) is shown.

**Figure S24:**
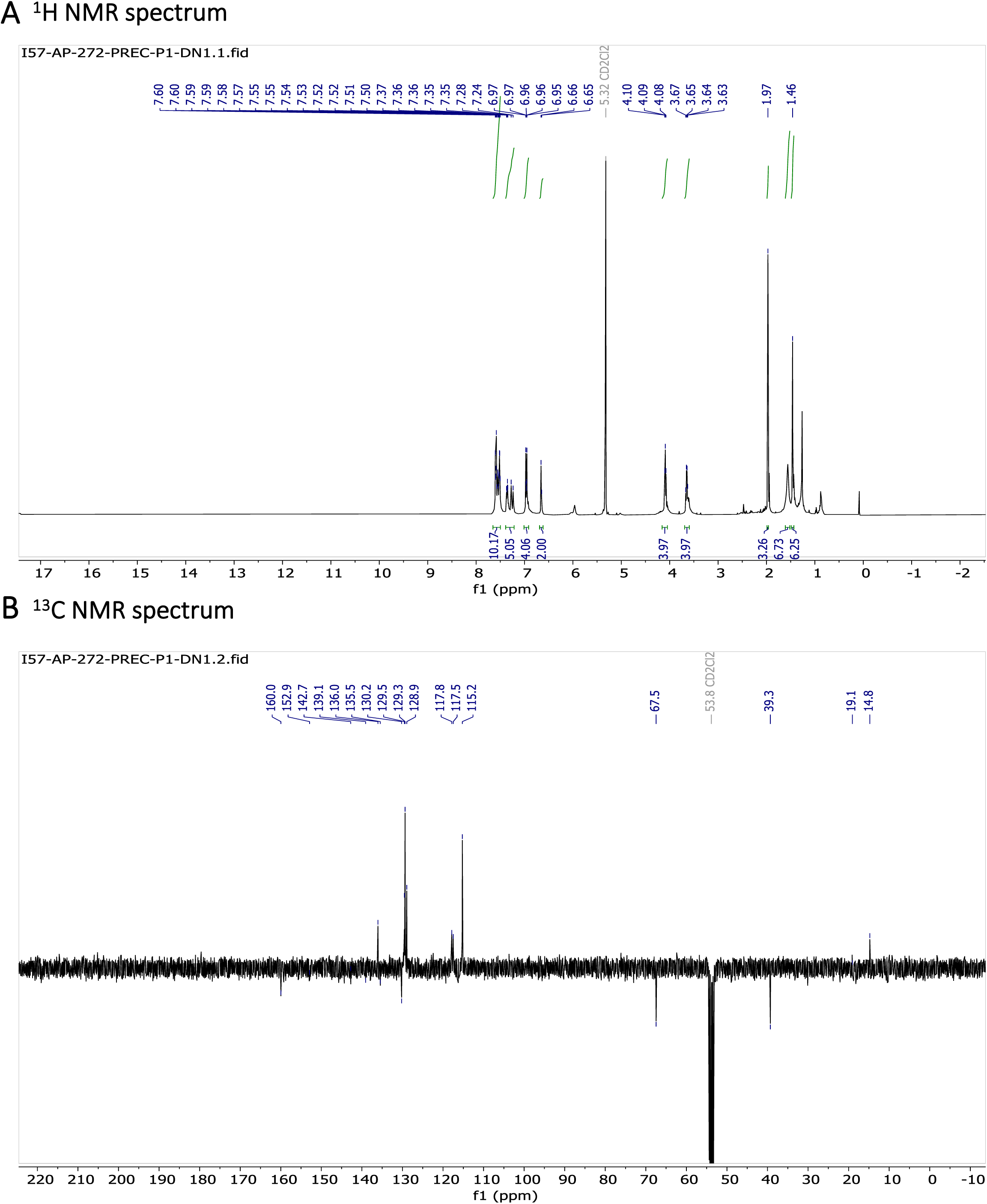
Characterization of BODIPY-NHAc. (A) ^1^H NMR spectrum and (B) ^13^C NMR spectrum are shown.

**Figure S25:**
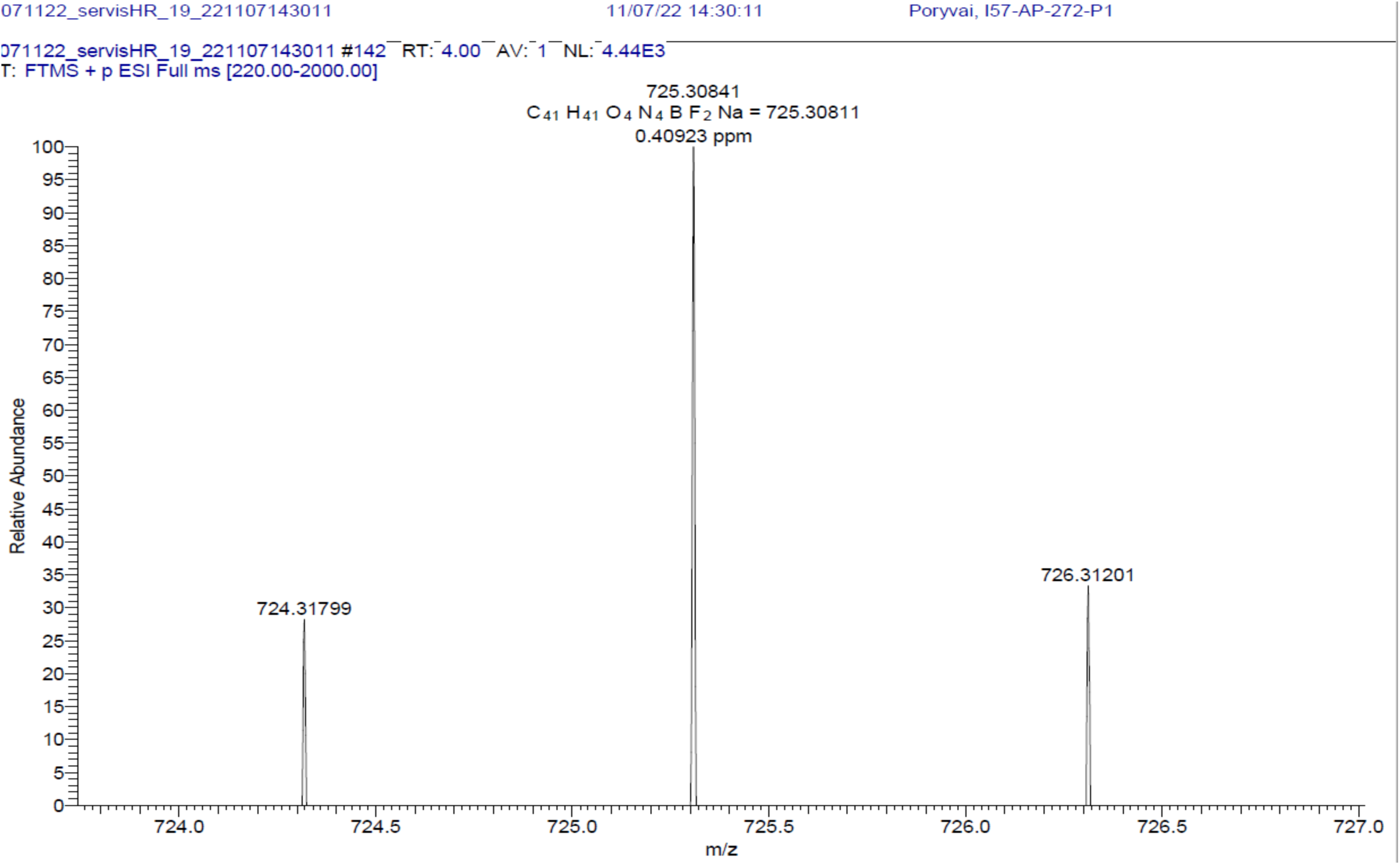
BODIPY-NHAc. HRMS spectrum (+EI) is shown

